# Light Activated BioID (LAB): an optically activated proximity labeling system to study protein-protein interactions

**DOI:** 10.1101/2022.10.22.513249

**Authors:** Omer Shafraz, Carolyn Marie Orduno Davis, Sanjeevi Sivasankar

## Abstract

Proximity labeling with genetically encoded enzymes is widely used to study protein-protein interactions in cells. However, the resolution and accuracy of proximity labeling methods are limited by a lack of control over the enzymatic labeling process. Here, we present a high spatial and temporal resolution technology that can be activated on demand using light, for high accuracy proximity labeling. Our system, called Light Activated BioID (LAB), is generated by fusing the two halves of the split-TurboID proximity labeling enzyme to the photodimeric proteins CRY2 and CIB1. Using live cell imaging, immunofluorescence, western blotting, and mass spectrometry, we show that upon exposure to blue light, CRY2 and CIB1 dimerize, reconstitute the split-TurboID enzyme, and biotinylate proximate proteins. Turning off the light halts the biotinylation reaction. We validate LAB in different cell types and demonstrate that it can identify known binding partners of proteins while reducing background labeling and false positives.

## Introduction

Protein-protein interactions (PPIs) are essential for cellular function, and decoding these interactions are crucial for understanding biological pathways in health and disease. While many methods have been developed to map protein interactomes (Qin et al., 2021), proximity labeling (PL) techniques are among the most widely used approaches, since they capture transient, dynamic PPIs in the near-native cellular environment.

PL employs enzymes such as promiscuous biotin ligases for proximity-dependent biotin identification (BioID) (Choi-Rhee et al., 2004; Kim et al., 2016; Ramanathan et al., 2018; Roux et al., 2012), peroxidases for peroxide-dependent biotin identification (APEX) (Martell et al., 2012), horseradish peroxidase (HRP) (Kotani et al., 2008), or Pup ligase PafA in PUP-IT (Liu et al., 2018). These enzymes are genetically fused to a protein of interest (the *bait*) and generate short-lived intermediate reactive biotins that covalently tag neighboring proteins (the *prey*). Other methods of tagging include using photosensitizers such as the enzyme miniSOG (Zhai et al., 2022) or photocatalyst-antibody conjugates to generate singlet oxygen (Müller et al., 2021) or activated carbenes (Geri et al., 2020) as alternate intermediates for targeting biotin-tagged probes. Subsequently, biotin-tagged prey proteins are enriched using streptavidin-coated beads and identified using mass spectrometry (MS).

BioID is preferred over other PL methods in many *in vitro* and *in vivo* applications, since peroxidases such as APEX and HRP require the addition of hydrogen peroxide and miniSOG produces reactive oxygen species, which can both be toxic to cells. However, BioID has long labeling times (>18 hrs) and requires an optimal temperature of 37° C (Roux et al., 2012). In an effort to improve BioID’s performance, TurboID has been introduced as a fast-labeling biotin ligase (10 min labeling time) that also retains activity at temperatures <37° C, which enables it to be used in organisms like flies, worms, yeast (Branon et al., 2018), and plants (Mair et al., 2019), as well as in cell culture (Shafraz et al., 2020). However, TurboID can biotinylate proteins using endogenous cellular biotin, which creates a large labeling background as well as cellular toxicity. To overcome this limitation, a ‘split-TurboID’ technology was recently introduced, which splits the TurboID enzyme into two inactive halves that only induce biotinylation when reconstituted by the addition of a chemical cofactor, rapamycin. While this reduces background labeling and increases spatial specificity (Cho et al., 2020a), the requirement that both halves of Split-TurboID be expressed in close proximity creates a level of background biotinylation unrelated to the addition of the cofactor (Cho et al., 2020b). Additionally, usage of a cofactor introduces the possibility of uncontrolled effects on cellular processes, and the time resolution is limited by the cofactor’s ability to diffuse to the cell compartment of interest. Furthermore, the difficulty of removing the factors from the media makes it difficult to control the cessation of the biotinylation reaction (Klewer and Wu, 2019).

These limitations in conventional proximity labeling can be overcome by developing new classes of proximity labeling technologies that can be activated on demand without the addition of chemical cofactors, and with high spatial and temporal control. We therefore designed a proximity labeling technology that is precisely triggered using blue (488 nm) light. Our system, called Light Activated BioID (LAB), is generated by fusing the two halves of the split-TurboID enzyme to genetically encoded *Arabidopsis thaliana* photodimeric proteins cryptochrome 2 (CRY2) and cryptochrome-interacting basic-helix-loop-helix (CIB1) (Liu et al., 2008). Upon exposure to blue light, CRY2 and CIB1 dimerize within 300 ms (Kennedy et al., 2010; Taslimi et al., 2016) and reconstitute the two halves of the split-TurboID enzyme, thereby inducing biotinylation. Since the photodimers have a half-life of ∼6 minutes (Kennedy et al., 2010; Taslimi et al., 2016), dissociation of CRY2 and CIB1 re-splits the TurboID and halts biotinylation. Since LAB dissociates within minutes after the blue light is turned off, background biotinylation is reduced and false positives as well as cellular toxicity due to over-biotinylation are minimized.

Here, as a proof of concept, we use live cell imaging, immunofluorescence, western blots, and mass spectrometry (MS) to show in Human Embryonic Kidney (HEK) 293T cells that LAB dimerizes and biotinylates proximal proteins when exposed to blue light. Next, by fusing LAB to the ubiquitous cell-cell adhesion protein E-cadherin (Ecad) in Madin-Darby canine kidney (MDCK) cells, we demonstrate that LAB can map Ecad’s known binding partners in a light-dependent manner, with higher accuracy and significantly fewer false positives compared to stand-alone TurboID.

## Results

### LAB dimerizes upon blue light illumination

We developed genetically encodable LAB constructs by fusing previously reported low-affinity split-TurboID (L73/G74) (Cho et al., 2020a) onto light-inducible dimerizing plant proteins CRY2 and CIB1. The N-terminal fragment of split-TurboID (spTN) and CIB1 were fused onto plasma membrane-targeted enhanced Green Fluorescent Protein (pmEGFP). The C-terminal split TurboID fragment (spTC) and CRY2 were fused onto mCherry and expressed in the cytoplasm (Fig. 1a). We hypothesized that, upon blue light exposure, CRY2-spTC-mCherry (*henceforth referred to as CryC*) would translocate to the plasma membrane and bind to CIB1-spTN-pmEGFP (*henceforth referred to as CibN*) (Fig. 1a).

**Figure 1:**
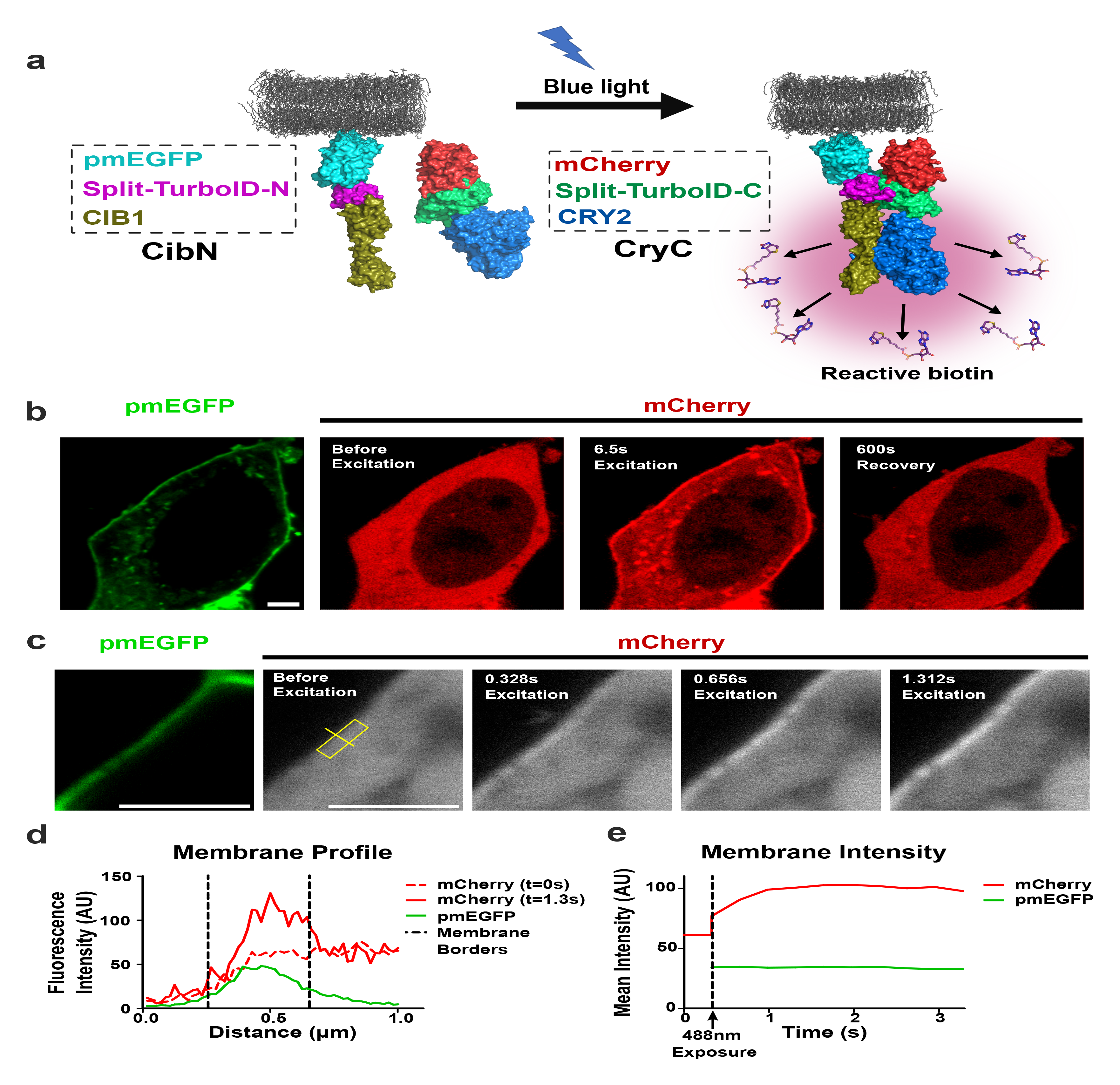
Light Activated BioID (LAB) dimerizes upon blue light illumination. (a) Schematic of LAB construct used in HEK293T cells. Fragment of Split-TurboID (N) and *Arabidopsis* protein CIB1 were fused onto prenylated plasma membrane targeted Enhanced Green Fluorescent Protein (pmEGFP) that localizes to the plasma membrane (CibN). The complementary Split-TurboID (C) fragment was fused to blue-light receptor protein Cryptochrome 2 (CRY2) and mCherry fluorescent protein (CryC) and expressed in the cytoplasm. When exposed to blue light, CRY2 interacts with CIB1 and reconstitutes the split-TurboID, which generates reactive biotin for proximity labeling. Both constructs were transiently co-expressed in HEK293T cells. (b) Confocal images show CibN localizing to the membrane and CryC diffused in the cytoplasm of the cells. When excited with a 488 nm laser, CryC localized to the membrane in seconds and dissociated completely in about 600 sec. Scale bar 3 µm. (c) A region of the membrane was excited with 488 nm laser and CryC accumulation was monitored. CryC accumulated at the membrane region in sub-seconds. Scale bar 3 µm. (d) CryC and CibN intensity was measured across the yellow line in *(c)* with vertical black dashed lines delineating the membrane on the graph. The fluorescence intensity measurements showed a strong correlation between CryC and CibN after light exposure. (e) Integrated CryC (red) and CibN (green) intensity was measured in the yellow box in *(c)* over time, with the vertical black dashed line indicating 488 nm laser excitation. CryC fluorescence reached the maximum intensity within seconds, while CibN fluorescence remained constant.

We transiently expressed both constructs in HEK293T cells and observed that CryC was distributed throughout the cytoplasm while CibN was localized at the plasma membrane (Fig. 1b). When illuminated with a 488 nm laser, CryC translocated to the plasma membrane within seconds of laser exposure; the residual mCherry signal in the cytoplasm even after laser exposure arose because CryC was expressed at high enough levels to saturate CibN on the membrane (Fig. 1b). When the laser beam was turned off, CryC dissociated back to the cytoplasm in ∼10 mins (Fig. 1b). These kinetics show that the fused constructs don’t alter the previously reported CRY2/CIB1 kinetics (Kennedy et al., 2010). Furthermore, when a selected region of the plasma membrane was illuminated with a 488 nm laser, CryC began accumulating in under a second and reached maximum accumulation in seconds (Fig. 1 c, d, e). This demonstrates that LAB can be activated in selected regions of the cell and split-TurboID will be bound for ∼ 10 mins from the end of illumination, biotinylating neighboring proteins and giving more temporal and spatial control over previous PL systems.

### LAB biotinylates only in the presence of blue light and biotin

Since split-TurboID shows significant biotin labeling after 1hr (Cho et al., 2020a), we transiently co-expressed CibN and CryC in HEK cells and exposed the cells to alternating 10-minute cycles of blue light (470 nm) and darkness for 1hr using a blue LED light in the presence of 100 µM biotin (Fig. 2a). We then fixed the cells and stained them for GFP, mCherry, and biotin using anti-GFP and anti-mCherry antibodies and Alexa-647 conjugated streptavidin (Sta) (Fig. 2 b-g, S1-6). Next, we measured the level of CibN, CryC, and Sta on the membrane; fluorescence intensities were normalized to CibN fluorescence intensity to account for differences in CibN expression levels in different cells (Fig. 2h, table S1). We also quantified the colocalization of CryC and Sta with membrane bound CibN (Fig. 2i, table S2). All p-values are catalogued in the supplementary information (tables S3, S4).

**Figure 2:**
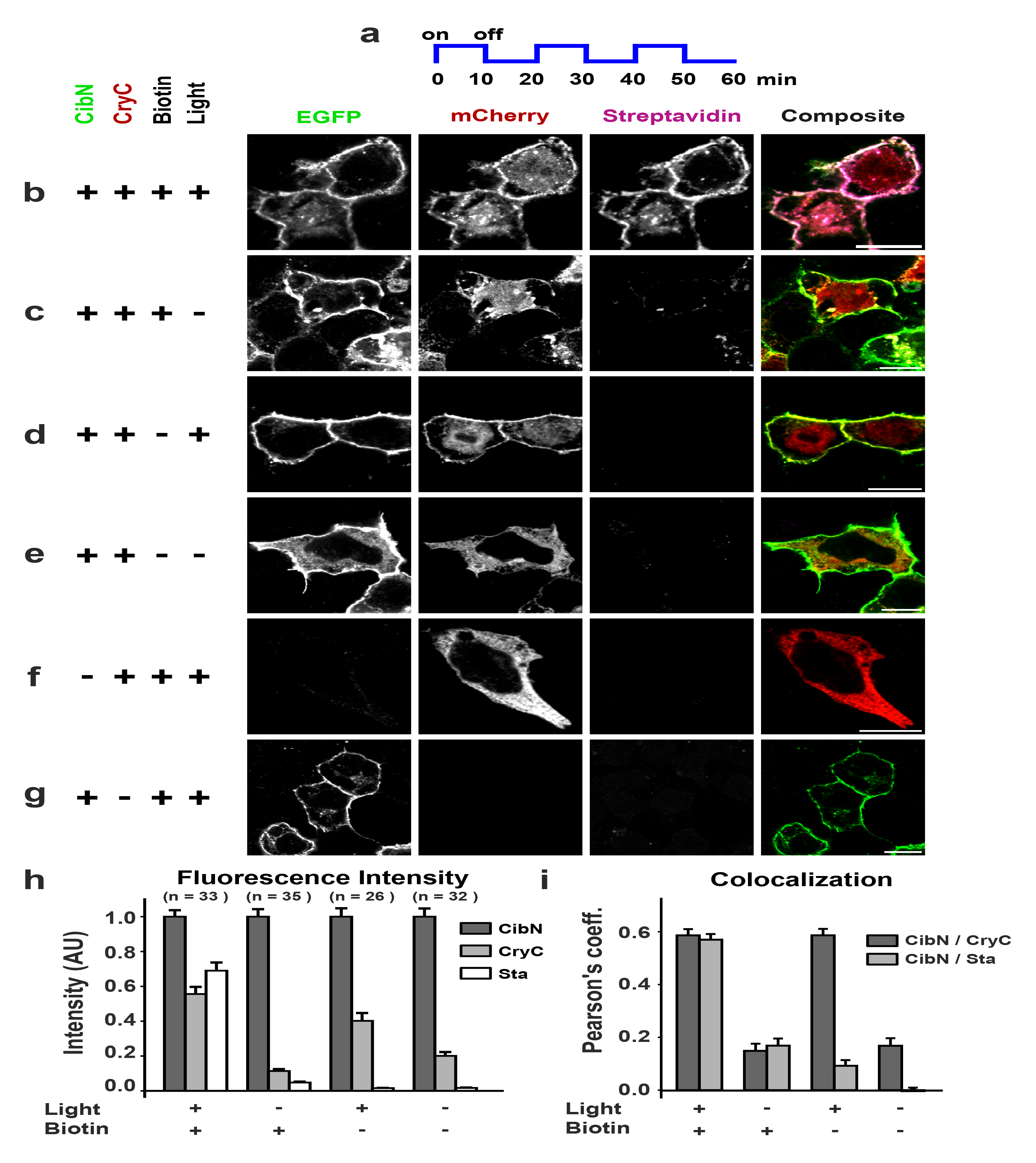
LAB biotinylation is light-dependent. CibN and CryC were transiently co-expressed in HEK 293T cells and, in the presence of 100 µM biotin, were exposed to 10 min cycles of alternating blue light and darkness for 1hr. The cells were fixed and stained for EGFP, mCherry, and biotin. (a) Schematic of light exposure sequence used. (b) Immunofluorescence showed that in the presence (+) of exogenous biotin and light, membrane-localized CibN (EGFP) and CryC (mCherry) dimerized and biotinylated proteins (Streptavidin, Sta) in the vicinity of the plasma membrane, with composite images confirming the colocalization of CibN, CryC, and Sta. (c) In the absence (-) of blue light, (d) in the absence of added biotin or (e) in the absence of both biotin and light, no biotinylation was observed. When only (f) CryC or (g) CibN were expressed, no biotinylation was detected, showing CibN and CryC dimerization are essential for proximity labeling. To facilitate an unbiased visual comparison of fluorescence intensities in (*b - g*), the panels in each column display identical minimum and maximum intensities. (h) Bar graph of CryC, CibN, and Sta fluorescent intensity averaged across number of cells from three biological replicates show significant biotinylation occurring only in the presence of light and biotin. CryC intensity proximal to membrane is significant only in the light positive conditions, showing it localizes with the membrane only in the presence of light. Fluorescent intensity of CryC and Sta proximal to membrane region was normalized to CibN to account for variation in CibN expression levels in different cells. Normalized fluorescence intensities are tabulated in table S1 while corresponding p-values are tabulated in table S3. (i) Average Pearson’s coefficient shows CibN and CryC colocalize in the presence of light, while Sta colocalizes to CibN only in the presence of light and biotin. Average Pearson’s coefficient values are tabulated in table S2 while corresponding p-values are tabulated in table S4. All scale bars 10 µm. Errors: s.e. n= number of cells from three biological replicates for all conditions.

In the presence of light and biotin, CryC showed a high colocalization with CibN at the plasma membrane, which resulted in high Sta fluorescent intensity corresponding to significant membrane proximal biotinylation (Figs. 2 b, h, i, S1, table S1, table S2). When samples were kept in darkness, the CryC did not localize to the membrane and no biotinylation was observed (Fig. 2 c, h, i, S2, table S1, table S2). Similarly, when cells were exposed to light in the absence of biotin, CryC had significant membrane fluorescence intensity and colocalization with CibN (Fig. 2 d, h, i, S3, table S2), but did not result in significant biotinylation, since Sta fluorescent intensity was very low at the membrane with negligible colocalization between Sta and CibN (Fig. 2 h, i, table S2). Furthermore, in the absence of both light and biotin, very low membrane CryC and Sta fluorescence intensity and CibN colocalization were measured corresponding to negligible biotinylation (Fig. 2 e, h, i, S4, table S1, table S2). Finally, no biotinylation was measured when only CryC (Fig. 2f, S5) or CibN (Fig. 2g, S6) was expressed. Hence, the immunofluorescence data demonstrate that CibN and CryC dimerize and biotinylate only when exposed to blue light.

### Ecad-LAB activity is light-dependent

Next, to validate LAB activity with a well-characterized protein and in a different cell line, we fused LAB to the ubiquitous, transmembrane protein E-cadherin (Ecad) in MDCK cells. Ecad is an essential cell-cell adhesion protein that plays key roles in the formation and maintenance of epithelial tissues and acts as a tumor suppressor (Xie et al., 2022). Besides binding homophilically, Ecad also has many recorded heterophilic transmembrane binding partners such as the desmosomal adhesion proteins Desmoglein-2 (Dsg2) and Desmocollin-3 (Dsc3) (Shafraz et al., 2020). We fused CIB1 and spTN onto Ecad-EGFP (ECibN) (Fig. 3a) and stabily expressed it with CryC in MDCK cells containing endogenous Ecad.

**Figure 3:**
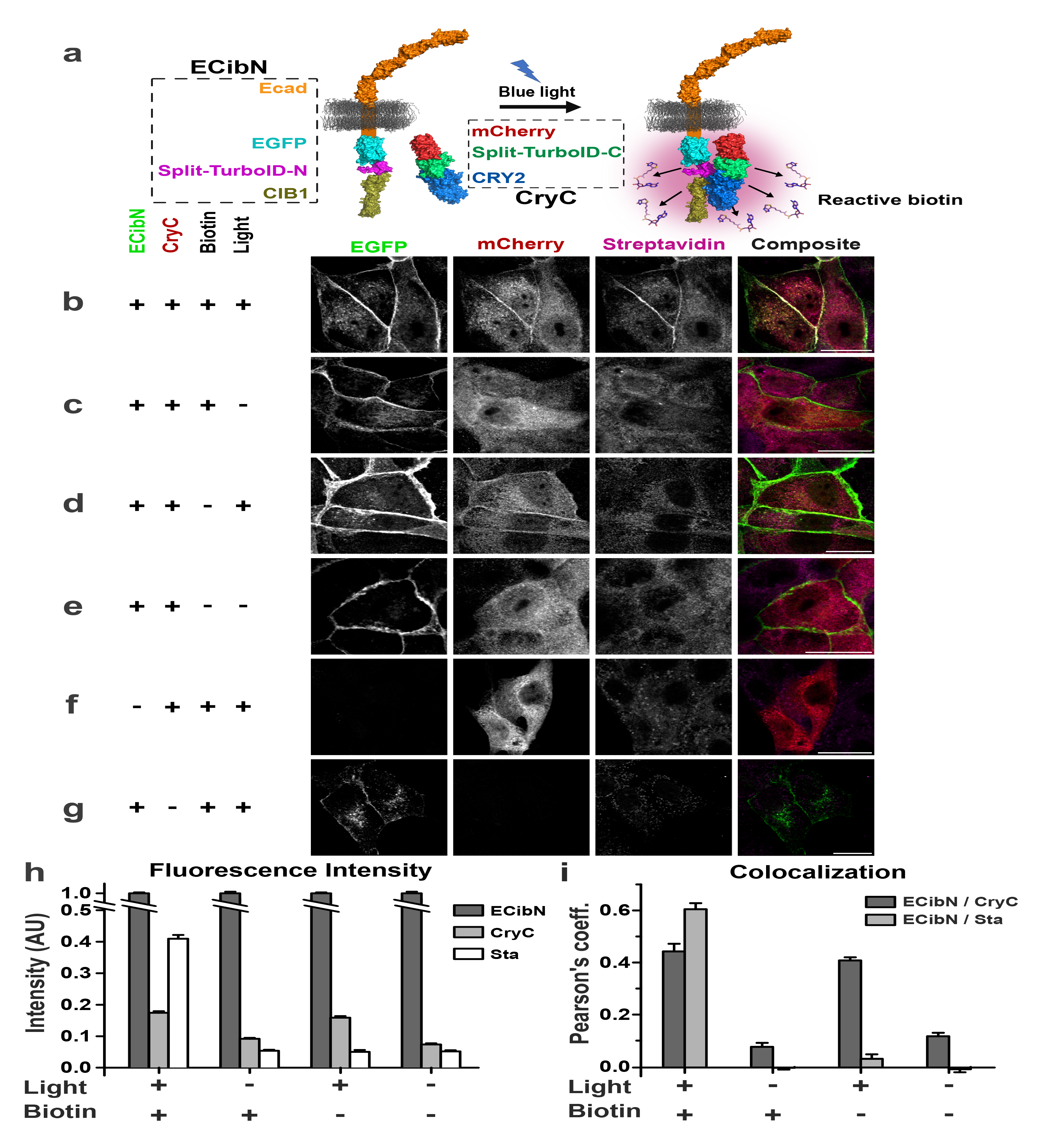
Ecad-CibN and CryC dimerize and biotinylate only when exposed to light. (a) Schematic of Ecad-LAB. Split-TurboID (N), and CIB1 were fused onto the C terminus of Ecad-EGFP to create ECibN. CryC was expressed in the cytoplasm. Both constructs were stably co-expressed in MDCK cells. When exposed to blue light, dimerization of ECibN and CryC reconstitutes the split-TurboID, which generates reactive biotin for proximity labeling. (b) Cells incubated with 100 µM biotin were illuminated with blue light, for an hour, on a 10 min on/off cycle. The cells were fixed and stained for EGFP, mCherry and biotin. ECibN localizes to the intercellular junction, and in the presence (+) of light and biotin dimerizes with CryC resulting in the biotinylation of proteins close to the membrane, which is confirmed with composite images showing the colocalization of CibN, CryC, and Sta. (c) In the absence (-) of light, (d) biotin or (e) both, no biotinylation is observed. When only (f) CryC or (g) ECibN were transiently expressed in MDCK cells, no biotinylation was observed. To facilitate an unbiased visual comparison of fluorescence intensities in (*b - g*), the panels in each column display identical minimum and maximum intensity. (h) Fluorescent intensity of CryC, ECibN, and Sta, normalized to ECibN intensity to account for variation in expression levels in different cells, shows that CryC only associates to the membrane in light and biotinylation is detected only in the presence of light and biotin. Normalized fluorescence intensities are tabulated in table S5 while corresponding p-values are tabulated in table S7. (i) Average Pearson’s coefficient shows ECibN and CryC colocalize in the presence of light, while Sta colocalizes to ECibN only in the presence of light and biotin. Average Pearson’s coefficient values are tabulated in table S6 while corresponding p-values are tabulated in table S8. All scale bars are 20 µm. Errors: s.e. n = 156 cells evenly split between three biological replicates for all conditions.

First, we confirmed that Ecad-CibN localizes to cell-cell junctions (Fig. 3b, S7-S12). Next, cells were exposed to alternating 10 min cycles of blue light and darkness for 1hr before fixing and immunostaining and CryC accumulation at the junction was monitored. To measure biotinylation and localization, ECibN, CryC, and Sta fluorescence intensity (normalized by ECibN fluorescence intensity to account for variation in ECibN expression levels) were measured. The high CryC and Sta fluorescence intensity near the membrane in the presence of light and biotin confirmed biotinylation activity, with high colocalization coefficients for ECibN with both CryC and Sta (Fig. 3 b, h, i, S7, table S5, S6). In the absence of light and the presence of exogenous biotin, CryC did not colocalize with ECibN, and a low membrane intensity and colocalization coefficient was measured (Fig. 3 c, h, i, S8, table S6). Additionally, no biotinylation was detected, with a negligible Sta membrane intensity and colocalization coefficient between ECibN and Sta (Fig. 3 c, h, i, S8, table S5, S6). This indicates that Ecad-LAB biotinylation is light dependent. Furthermore, when cells were illuminated with blue light in the absence of biotin, even though ECibN and CryC showed colocalization and CryC had an increased membrane intensity (Fig. 3 d, h, i, S9, table S6), the relative Sta intensity and colocalization was not significant (Fig. 3 d, h, i, table S5, S6). Similarly, in the absence of both light and biotin, ECibN/CryC did not colocalize or show a substantial CryC or Sta membrane signal, with no colocalization between ECibN and Sta (Fig. 3 e, h, i, S10, table S6). Finally, when only CryC (Fig. 3f, S11) or ECibN (Fig. 3g, S12) was expressed, there was no biotinylation observed. Thus, the immunofluorescence data for ECibN and CryC in MDCK cells confirm that LAB can be applied to other proteins of interest, in different cell lines, and that when exogenous biotin is provided, its biotinylation activity depends only on exposure to light. All p-values are catalogued in supplementary information (tables S7, S8). It is important to note that the presence of a high background in the Sta channel for MDCK cells was due to a low ECibN expression level and not due to off-target biotinylation since the background levels for the biotin positive and negative conditions were similar (Fig. 3 b-e).

Finally, to quantify how quickly Ecad-LAB biotinylates proximal proteins, we performed immunofluorescence analysis for MDCK cells exposed to light for different durations (1min, 10min, and 30min; Fig. S13). The data shows statistically significant biotinylation after just 1min of light exposure, indicating that Ecad-LAB is indeed biotinylating on this timescale (Fig. S13 a, d, e).

### Benchmarking LAB against stand-alone TurboID

To benchmark LAB performance, we compared Ecad proteomes generated using both LAB and stand-alone full-length TurboID. A biotinylated LAB proteome was generated using MDCK cells stably expressing Ecad-LAB. The cells were incubated in 100µM biotin and were either exposed to blue light for 1 hr using 10 min on/off cycles or were kept in darkness. To generate a corresponding biotinylated TurboID proteome, MDCK cells were stably transfected with Ecad conjugated to full-length TurboID (Ecad-Turbo). The Ecad-Turbo cells were either incubated with or without 100µM biotin for 1hr. The biotinylated proteins generated by Ecad-LAB and Ecad-Turbo were then captured from the corresponding cell lysate using streptavidin-coated magnetic beads. The captured proteins were trypsin digested before LC-MS/MS was performed for three replicates per condition (Ecad-LAB) or two replicates per condition (Ecad-Turbo).

The Ecad-LAB MS data showed significant enrichment of proteins in cells illuminated with blue light compared to those kept in darkness (Fig. 4a). We then compared Ecad-LAB hits in the presence and absence of light against Ecad-Turbo in the presence and absence of biotin (Fig. 4). This data demonstrated that Ecad-LAB had 300 positive hits and only 3 negative hits (Fig. 4a). In contrast, Ecad-Turbo had 247 positive hits and 139 negative hits (Fig. 4b). Well-established members of the core Ecad/catenin complex (Guo et al., 2014) such as Ecad, β-catenin, α-catenin, p120-catenin, and vinculin were highly enriched in Ecad-LAB. Similarly, well-established Ecad transmembrane binding partners such as Dsg2 and Dsc3 were prominent hits in Ecad-LAB (Shafraz et al., 2020) (Fig. 4a). Importantly, all the hits (positive candidates, negative candidates, and non-candidates) in Ecad-LAB were also hits in Ecad-Turbo, with the expected exceptions of the light-activated dimer components (Cry/CIB) (Fig. 4c). This demonstrates that Ecad-LAB is at least as accurate as Ecad-Turbo, with an increased precision resulting in fewer extraneous hits.

**Figure 4:**
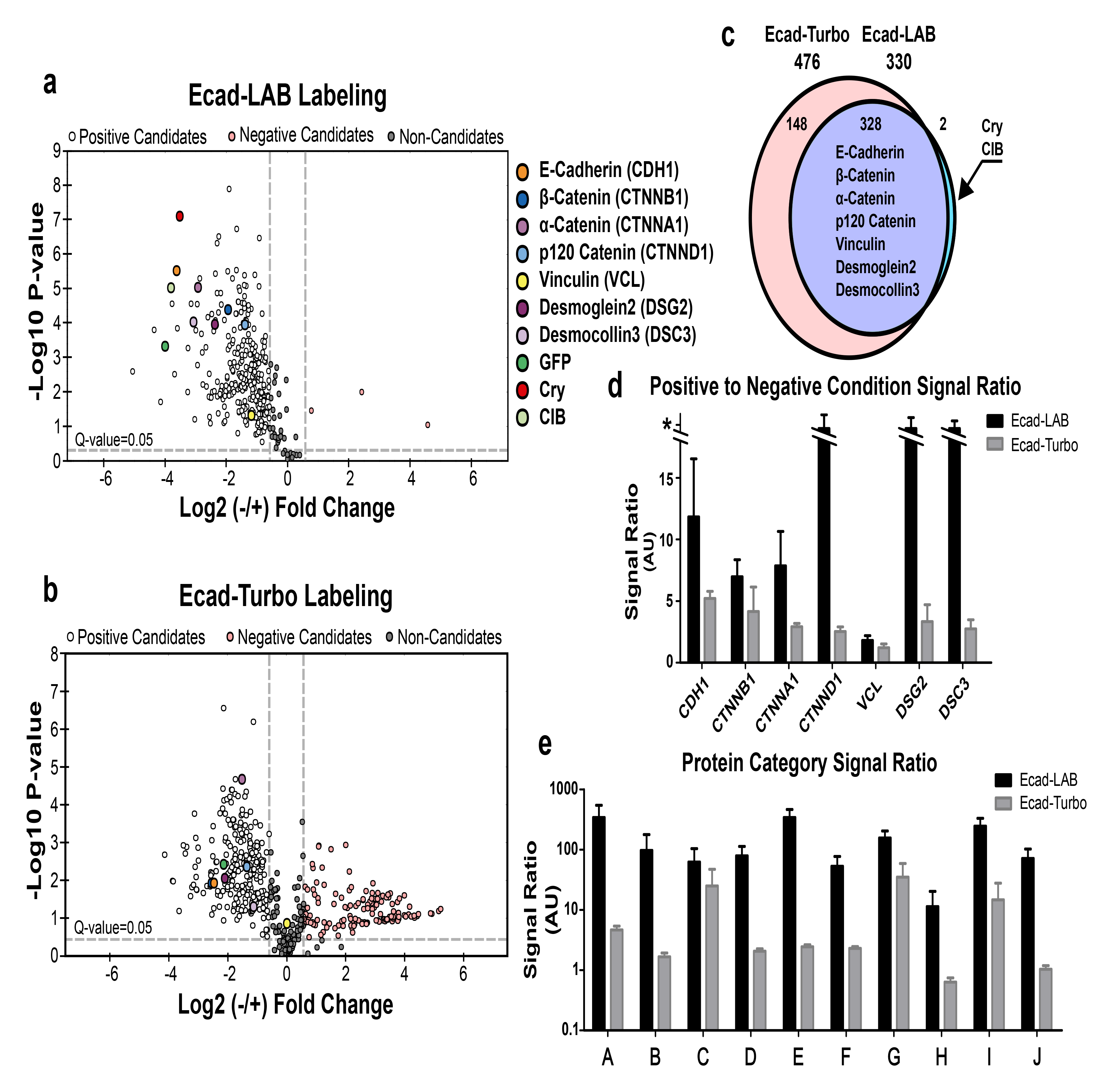
Mass spectrometry shows that Ecad-LAB is more accurate and precise than Ecad-Turbo. a) Volcano plot showing the enrichment of biotinylated proteins in Ecad-LAB, in the presence (positive; white) and absence (negative; pink) of light. Known Ecad-associated proteins are labeled in different colors and are highly enriched. b) Volcano plot showing the enrichment of biotinylated proteins in Ecad-Turbo, in the presence (positive; white) and absence (negative; pink) of biotin. There are many more negative “hits” in Ecad-Turbo compared to Ecad-LAB. c) Every hit (positive candidates, negative candidates, and non-candidates) in Ecad-LAB was also a hit in Ecad-Turbo, with the expected exceptions of Cry and CIB. d) Comparison of positive/negative signals for several key Ecad binding proteins highlighted in (*a*). Ecad-LAB has a universally higher signal ratio. Proteins without any signal in the negative condition were given an arbitrarily high ratio value (*) to avoid dividing by zero. e) Signal ratio for broad categories of proteins, graphed on a log scale. Categories were chosen based on previously published categories in the Ecad interactome (Shafraz et al., 2020), as follows: A: Adhesion Receptors; B: Cytoskeletal components and motors; C: Kinases and Phosphatases; D: Membrane binding adaptors, receptors, and transporters; E: Other/Unknown/Secreted; F: Actin and microtubule binding and dynamics-associated; G: Adaptors; H: Metabolic Enzymes; I: Chaperone, trafficking, degradation, and GTPase regulation; J: DNA and Translation.

Comparison of the ratio of positive/negative signal levels for Ecad and its key binding partners (highlighted in Fig. 4a) demonstrated that Ecad-LAB has a universally higher signal/background ratio compared to Ecad-Turbo (Fig. 4d). This trend extended for the entire Ecad-LAB interactome as seen in the signal/background ratios for broad categories of proteins (Fig. 4e). In this figure the Ecad-LAB interactome was sorted into previously published categories (Shafraz et al., 2020) as described in the figure legend. We further directly compared the biotin negative and light negative conditions of Ecad-Turbo and Ecad-LAB respectively, which showed an almost universally higher absolute protein level present in the negative Ecad-Turbo condition compared to the negative Ecad-LAB condition (Fig. S14). Taken together, this data quantitatively demonstrates that LAB has a significantly lower background biotinylation than conventional TurboID.

We also used MS to confirm LAB’s activity in HEK293T cells transiently co-expressing CibN and CryC. As described above for Ecad-LAB, the cells in the presence of biotin were either exposed to blue light or darkness and biotinylated proteins were isolated, trypsin digested and analyzed using LC-MS/MS (three replicates per condition). MS data showed significant enrichment of proteins in cells illuminated with blue light compared to those kept in darkness (Fig. S15a). Although membrane-bound GFP does not have any prey in HEK293T cells, CIB1 and GFP, which are close to the Split-TurboID, served as proxy prey and were significantly enriched (Fig. S15a). Finally, principal component analysis (PCA) showed two separate clusters for the light-exposed samples and the controls, implying distinctive variations between both groups (Fig. S15b).

### Benchmarking biotinylation efficiency of LAB against stand-alone TurboID

We used western blots to benchmark the biotinylation efficiency of Ecad-LAB exposed to light for different durations (1hr, 3hrs, 5hrs, 18hrs) against Ecad-Turbo incubated in biotin for 10 mins (Fig. S16). To prevent phototoxicity in Ecad-LAB cells due to their long exposure to blue-light, a shorter on/off light cycle (1min on: 5min off) was used. Western blots were performed on whole cell lysates with an equal amount of protein loaded in each lane; biotinylated proteins were stained with anti-biotin HRP.

While Ecad-Turbo incubated with biotin for 10 mins had higher biotinylation levels than Ecad-LAB at all measured timepoints (Fig. S16a), the larger number of false positives in Ecad-Turbo (Figs. 4, S14) made a direct comparison of biotinylation from western blot bands difficult. We therefore compared the biotinylation efficiency of Ecad-LAB for 1 time point (1 hr) against Ecad-Turbo and used a single band at ∼130kDa, which is present in the Ecad-LAB and Ecad-Turbo lanes (but not in the untransfected control), for subsequent analysis (Fig. S16a, S16b).

It is important to note that while every Ecad-Turbo protein is capable of biotinylating a target, Ecad-LAB has a Pearson’s correlation coefficient of 0.44 (see Fig. 3), implying that only ∼44% of ECibN proteins are bound to CryC and capable of biotinylating. Furthermore, differences in the expression levels of ECibN and Ecad-Turbo could also impact biotinylation efficiency. We therefore intensity corrected the Ecad-Turbo and Ecad-LAB lanes for their differing expression levels (determined using an Ecad western blot, Fig. S16c) and for ECibN/CryC colocalization (using experimentally determined Pearson’s coefficients, Fig. 3). The intensity corrected bands were used to directly compare the relative efficiencies of Ecad-Turbo and Ecad-LAB (Fig. S16 d-f). The ratio between the positive and negative condition bands for each construct showed that intensity corrected Ecad-LAB with 1 hour light exposure had a similar biotinylation efficiency as Ecad-Turbo incubated with biotin for 10 minutes (Fig. S16f). In contrast, stand-alone Split-TurboID (albeit in non-correlation corrected data) needed 4 hours to approach 1 minute of full-length TurboID labeling (Cho et al., 2020b).

To compare biotinylation efficiency of LAB and stand-alone TurboID in HEK cells, we transiently expressed a full TurboID construct (CIB1-TurboID-pmEGFP) and incubated in 100 µM biotin for 1 hr. We also transiently co-expressed CryC with CibN in HEK cells and exposed these cells to blue light in the presence of biotin for 10 min on/off intervals for 1 hr. Both cells were then lysed and biotinylated proteins were captured using streptavidin-conjugated beads and used for western blots, with the eluted protein stained with Sta-HRP (Fig. S17). Western blots showed significantly higher biotinylation in light-exposed HEK293T cells compared to cells kept in darkness. However, the biotinylation efficiency of LAB was less than cells expressing TurboID, likely due to the lower activity and background from LAB (Fig. S17a).

While we did not have any known prey for CibN in HEK293T cells, we expected the GFP and mCherry portions of CibN and CryC to be biotinylated, as these would be directly proximal to the reconstituted Split-TurboID. When we stained eluted protein with GFP and mCherry antibodies, we found a much higher presence of these proteins in the light-positive condition than the light negative condition, with no staining occurring on a blank control (Fig. S17 b, c). This confirms that LAB biotinylates proximal proteins in a light-dependent manner and identifies expected binding partners.

## Discussion

For LAB, we used split-TurboID as the labeling enzyme due to its well characterized biotinylation kinetics. Similarly, we used the photodimer pair CRY2 and CIB1 because they are one of the most widely-used photodimer systems, and have been tested in many different cell lines as well as in living organisms to investigate a wide variety of cellular functions (Taslimi et al., 2016). Importantly, CRY2/CIB1 binding kinetics complement those of Split-TurboID by remaining associated long enough after light exposure to allow for sufficient biotinylation, while not remaining bound too long post-exposure so that the labeling reaction can be arrested with the removal of light.

Using immunofluorescence, western blots, and MS, in both HEK293T and MDCK cells, we demonstrated that LAB can be selectively activated through researcher-controlled biotin and light application. We were able to demonstrate statistically significant biotinylation after only one minute of light exposure, showing that functional complementation of LAB is achieved on a rapid timescale (Fig. S13a). LAB’s specificity was validated through MS analysis showing light exposure-dependent enrichment of many proteins known to bind to Ecad, including the α/β- catenin/vinculin complex that links Ecad to the actin cytoskeleton (Fig. 4a). Importantly, these and other known Ecad binding partners were highly enriched in the light positive condition compared to the light negative condition, with a signal to background ratio an order of magnitude higher than that of Ecad-conjugated full-length TurboID under biotin positive and negative conditions (Fig. 4e). Additionally, not every major Ecad binding partner was successfully detected as a positive candidate by Ecad-Turbo, possibly a result of its high background labeling reducing data resolution (Fig. 4b). Taken together, this shows that LAB successfully labels proximal proteins in a light-dependent manner with higher specificity than full-length TurboID.

Since LAB uses split-TurboID and light for biotinylation, it is reasonable to expect that LAB will efficiently biotinylate target proteins in different cellular compartments. It has already been demonstrated that both split-TurboID (Cho et al., 2020b) and stand-alone TurboID (Branon et al., 2018) are capable of biotinylating target proteins in different organelles with distinct pH, redox environments, and endogenous nucleophile concentrations. Additionally, since LAB relies on light rather than a diffusion-limited chemical cofactor, it can easily be activated in different cellular compartments. The main consideration will be expressing LAB sufficiently in different compartments, which is more a function of construct and bait design than of its biotinylation ability. Importantly, truncated versions of the CRY2 and CIB1 proteins have been developed (Taslimi et al., 2016), that can be used in LAB if large constructs would affect expression and localization of bait proteins tagged with LAB.

It has previously been shown that CRY2 can homodimerize when illuminated with blue light (Wang and Lin, 2020). However, this is not an issue for LAB since our construct only has the inactive C-terminus of Split-TurboID associated with CRY2. Consequently, CRY2 homodimerization cannot reconstitute a functional TurboID. This is supported by our data showing that no biotinylation is observed in the CryC-only condition (Fig. 2f). Furthermore, previous studies show that CRY2 homodimerization does not prevent interaction with CIB1 (Bugaj et al., 2013).

A consideration when using LAB is the possibility of phototoxicity after long-duration exposure to the activating blue light source. Due to Cry/CIB’s binding half-life of approximately 5 minutes, alternating 10- minute cycles of light and darkness allowed significant dimerization and biotinylation, while reducing phototoxicity for our shorter-term (< 1 hr) experiments. For longer duration experiments (see Fig. S16), we used a shorter light cycle (1min on, 5min off) to further reduce phototoxic effects while still maintaining maximal Ecad-LAB activation. However, for even longer-term experiments, the optimal light intensity and exposure cycle will likely need to be determined based on the cells and application.

Recently, a different opto-dimerization system, iLID, has been used with a new split-TurboID variant (Chen et al., 2022). However, iLID has a half-life of under a minute (Kennedy et al., 2010; Taslimi et al., 2016), which may not provide enough time for appreciable labeling to occur before the dimers unbind and inactivate Split-TurboID. Furthermore, in this opto-dimerization study, TurboID was split at a previously uncharacterized location (G99/E100) and consequently, the rate of biotinylation of this new split-TurboID variant is unknown (Chen et al., 2022).

Another PL tool using light-activated miniSOG has been recently developed (PDPL), which uses singlet oxygen to create electrophilic residues on prey proteins, which can be subsequently bound to an alkyne chemical probe and pulled down using click chemistry (Zhai et al., 2022). However, the aniline probe used with PDPL can be toxic to cells (Wang et al., 2016; Zhai et al., 2022). Furthermore, miniSOG has a labeling radius of up to 70nm, which is much larger than TurboID’s 10nm labeling radius (Kim et al., 2014). While a larger labeling radius is appropriate for identifying compartmentalization of proteins, LAB’s smaller labeling radius is more suited for discovering direct PPIs (Cho et al., 2020b; Zhai et al., 2022). Another light-activated PL tool, MicroMap, which uses an antibody conjugated to an iridium photocatalyst to label nearby proteins via carbene intermediates, has also recently been developed (Geri et al., 2020; Seath et al., 2021). However, the necessity of a primary antibody that binds to the bait protein at a convenient location limits MicroMap’s applications.

Unlike CRY2 and CIB1 which photodimerize in the presence of blue light, photodimerization activated by longer wavelength light could be more effective for PPI identification in deeper tissue. However, the two known red light activated photo-hetrodimers (PhyB/PIF3 and PhyB/PIF6), require light exposure at different wavelengths for both activation and inactivation (Klewer and Wu, 2019). We therefore chose a photodimer pair that dissociated in the dark, rather than through alternate wavelength exposure, for ease of use. Importantly, since CRY2/CIB1 dimerization can be triggered using two-photon excitation (Kennedy et al., 2010), this may allow LAB to map PPIs in deeper tissue *in vivo*.

We anticipate that the ability to activate LAB using light, coupled with its high temporal resolution, can be exploited to interrogate differences in PPIs at different time points in the cell cycle. We also expect that by using focused activating light, LAB can be used to identify differences in PPIs at distinct cellular locations. Finally, like TurboID, which has previously been used to map extracellular PPIs (Shafraz et al., 2020), we anticipate that LAB can also be used to investigate extracellular protein interactions with high spatial and temporal resolution.

## Materials and Methods

### Cloning of plasmid constructs

CibN was generated by restriction digesting a CIB-pmEGFP plasmid (Addgene plasmid # 28240, (Kennedy et al., 2010)) at AgeI and NheI sites and fusing PCR amplified CIB1 and spTN (Addgene plasmid # 153002, (Cho et al., 2020a)) using Gibson assembly. SpTN was connected to CIB1 and pmEGFP with linkers V5-KGSGSTSGSGTG (Linker 1) and GSGPVAT. CryC was constructed by inserting PCR amplified spTC (Addgene plasmid #153003, (Cho et al., 2020a)) onto restriction-digested Cry2-mCherry (Addgene plasmid # 26871, (Kennedy et al., 2010)) at XmaI sites with linkers ARGKGSGSTSGSG and KGSGDPPVAT.

To design ECibN, V5-spTN-CIB-pmEGFP was constructed by restriction digesting CIB1- pmEGFP with NheI and inserting PCR amplified Linker1-spTN-KGSGAT. Then, Ecad-EGFP plasmid was restriction digested at NotI and XhoI sites and EGFP was reintroduced without the stop codon. Next, NotI and HindIII sites of Ecad-EGFP were restriction digested and PCR amplified V5-spTN-CIB was inserted using Gibson assembly. Ecad-EGFP plasmid was a kind gift from Prof. Soichiro Yamada at the University of California, Davis. All enzymes were high efficiency enzymes from New England Biolabs. All primers used are listed in SI (table S9).

For development of a dual-expressing stable Ecad-LAB cell line, a Blasticidin (BSD) resistant CryC plasmid was generated by replacing spTN from Addgene plasmid #153002 (Cho et al., 2020a) using BstBI and NheI enzyme digestion and inserting CryC using Gibson assembly.

### Cell culture and transfection

HEK293T cells were grown in High Glucose (4.5 g/L) Dulbecco’s modified Eagle’s medium (DMEM, Gibco) cell culture media with 10% fetal bovine serum (Gibco) and 1% Penicillin-Streptomycin (PSK) (10,000 U/mL, Life Technologies). MDCK cells were cultured in Low Glucose (1 g/L) DMEM (Gibco) cell culture media with 10% fetal bovine serum and 1% PSK.

HEK cells were transiently transfected at 80% confluency with CibN and CryC plasmids using polyethylenimine (PEI) (Polysciences, Inc.). Plasmids and PEI were diluted in Opti-MEM (Gibco) at a 1µg:5µg DNA:PEI ratio. After five minutes, both solutions were mixed and incubated for 90 min and then added to the cells. In order to develop the MDCK stable lines, cells were transfected with the ECibN and BSD-CryC plasmids at 30% confluency using lipofectamine 3000 (Invitrogen) with the same DNA ratio. After 24 hours the cells were passaged and sparsely seeded onto p150 dishes. After allowing a further 24 hours for cell attachment and construct expression, 500 µg/mL G418 and 5 µg/mL BSD were added to the media for antibiotic selection. Colonies were picked after they reached ∼2 mm in size and transferred to a 96 well plate. Colonies were fluorescently imaged to detect expression, and top co-expressing colonies were further expanded. Expanded colonies were further imaged and the brightest clone with the most uniform expression was chosen for experiments.

### Live cell imaging

HEK cells were imaged >12 h post-transfection using a 561 nm laser to visualize mCherry and a 488 nm laser to image EGFP and to photo-excite CryC. Images were reconstructed using ImageJ. For the intensity profile of CryC accumulation, a 1µm line was drawn across a selected membrane region and the raw pixel intensity across the line was plotted (Fig. 1c). Measurements were taken using ImageJ’s Plot Profile analysis feature. To plot CryC accumulation over time, a 0.225µm by 0.900µm box was drawn on the cell membrane bracketing the intensity profile line (Fig. 1c). Measurements for each frame were taken using ImageJ’s ROI mean intensity measurement capability.

### Immunofluorescence

Cells were transferred to low glucose, phenol red-free DMEM (Gibco) with 10% FBS, incubated with 100 µM biotin and exposed to blue light in 10 min intervals for 1 hr using a blue LED light source (Blue box Pro Transilluminator, miniPCR bio). Control cells were kept in dark. Then, cells were immediately fixed using 3% paraformaldehyde and 0.3% Triton X-100 in phosphate-buffered saline (PBS) for 10 min and blocked with 1% bovine serum albumin (BSA) and 0.3% Triton X-100 in PBS for 30 min. Anti-GFP antibody (rabbit polyclonal, Rockland Immunochemicals) (HEK293T cells), Anti-GFP antibody (chicken polyclonal, Rockland Immunochemicals) (MDCK cells), Alexa Fluor 488-conjugated Goat anti-rabbit antibody (Life Technologies) (HEK293T cells), and Alexa Fluor 488-conjugated Goat anti-chicken antibody (Invitrogen) (MDCK cells) were used to detect GFP-tagged proteins. Anti-mCherry (mouse monoclonal, Invitrogen) (HEK293T cells), Anti-mCherry (rabbit monoclonal, Invitrogen) (MDCK cells), Alexa Fluor 568-conjugated Goat anti-mouse antibody (Invitrogen) (HEK293T cells), and Alexa Fluor 568-conjugated Goat anti-rabbit (Invitrogen) (MDCK cells) were used to detect mCherry. Biotinylated proteins were identified with Alexa Fluor 647-conjugated streptavidin (Invitrogen). Primary antibodies were incubated for 1 hr (1:1000 dilution in PBS with 1% BSA and 0.3% Triton X-100) and secondary antibodies were incubated for 30 min (1:1000 dilution in PBS with 1% BSA and 0.3% Triton X-100). Cells were imaged using a Leica Microsystems Stellaris 5 confocal setup with 63x/1.40 NA oil objective. Images were reconstructed using ImageJ.

### Image analysis

All analysis was done in ImageJ. In order to reduce background and oversaturated pixel contribution, all images were processed with a Gaussian filter with a sigma of 11 (HEK images) or 15 (MDCK images). The filtered image was then subtracted from the original image. CibN/ECibN expression was used to determine the area to be measured, and a threshold mask was created using the CIB-GFP signal using a threshold minimum of 50 on the Threshold tool. Membrane ROI’s were hand-chosen with the wand tool from the CibN/ECibN GFP channel using this mask. Since CibN and ECibN localize to the cell membrane, the ROIs only contain the plasma membrane. Higher Gaussian sigmas were used on MDCK cells due to the lower expression of ECibN. The ROIs were used for both fluorescence intensity measurements and colocalization analysis. Colocalization analysis was carried out on the processed images using the Coloc2 plugin with Costes threshold regression, a PSF of 3, and 10 Costes randomizations. Unthresholded Pearson’s coefficients were used from the outputs. Intensity measurements for all channels were taken on the processed images using the Multi Measure functionality on the ROI manager.

### Sample preparation for MS and Western Blot analysis

MDCK cells stably expressing Ecad-LAB or Ecad-Turbo and HEK cells transiently transfected with LAB were cultured on p150 dishes. The cells were incubated with 100 μM biotin and exposed to blue light on a 10 min on/off cycle for 1 hr (LAB, Ecad-LAB) or exposed to 100 μM biotin alone (Ecad-Turbo). Control cells were kept in darkness (LAB, Ecad-LAB) or without exogenous biotin (Ecad-Turbo). After incubation, cells were washed three times with PBS, scraped, and centrifuged. Pelleted MDCK cells were resuspended in M2 lysis buffer (50 mM Tris pH 7.5, 150 mM NaCl, 1% sodium dodecyl sulfate (SDS), 1% Triton X-100 (Muinao et al., 2018)) with 5 μL/mL of protease inhibitor mixture (Sigma-Aldrich), and 1 μL/mL Benzonase Nuclease (250 U/μL) (Millipore-Sigma). Pelleted HEK293T cells were resuspended in lysis buffer (50 mM Tris pH 7.5, 150 mM NaCl, 0.4% sodium dodecyl sulfate (SDS), 1% octyl phenoxy poly(ethyleneoxy) ethanol (IGEPAL CA-630), 1.5 mM MgCl_2_, 1 mM EGTA) with 2 μL/mL of protease inhibitor mixture (Sigma-Aldrich), and 1 μL/mL Benzonase Nuclease (250 U/μL) (Millipore-Sigma). The lysate was flash frozen in dry ice and quickly thawed at 37 °C. The lysate was then incubated for 30 min at 4 °C and sonicated at 30% duty ratio for 1 min. Next, the lysate was centrifuged at 13,000 rpm for 30 min at 4 °C. After that, the concentrations of the supernatant were measured with an RC DC protein assay kit (Bio-Rad). Supernatant concentration was adjusted to the lowest concentration and lysis buffer was added to bring all volumes to 1mL. Supernatant was incubated with 50 μL (HEK293T) or 100 μL (MDCK) of superparamagnetic streptavidin conjugated beads (Dynabeads MyOne Streptavidin C1, Invitrogen) and rotated overnight at 4 °C. The next day, beads were washed with the lysis buffer. Then, beads were washed with the 2% SDS in 50 mM Tris, pH 7.4, and twice with lysis buffer.

For MS sample preparation, magnetic beads loaded with biotinylated proteins were washed three times with 50 mM ammonium bicarbonate (NH_4_HCO_3_) and resuspended in 60 μL of 50 mM NH_4_HCO_3_ containing 5 µg of trypsin for overnight digestion at 37 °C in a shaker. The resulting peptides were recovered from the beads by removing the supernatant after pelleting the beads via magnetic holder, and tryptic peptides from the supernatant were dried in a vacuum centrifuge and reconstituted in 0.1% formic acid. Tryptic peptides were analyzed using nano-scale liquid chromatographic tandem mass spectrometry (nLC-MS/MS).

For MDCK western blots, whole cell lysate was used, with an identical amount of total protein loaded in each lane, determined using DC assay concentration measurements. Lysate was run on a 4-15% Mini-Protean TGX Precast Protein Gel (Bio-Rad) before being transferred to a nitrocellulose membrane and stained for biotin using HRP-conjugated Anti-biotin (1:1000 dilution, Cell Signaling Technology). WesternBright ECL HRP substrate (Advansta) was used to detect the protein.

For the HEK293T western blots, after capturing the biotinylated proteins, beads were incubated with 60 µl of 2.5 mM biotin at 95 °C for 5 min. Then, the eluted biotinylated proteins were collected and run on a 4-15% Mini-Protean TGX Precast Protein Gel (Bio-Rad) before being transferred to a nitrocellulose membrane and stained for biotin, GFP, and mCherry using Sta conjugated with Horseradish Peroxidase (HRP) (1:1000 dilution, Invitrogen), Anti-GFP antibody (rabbit polyclonal, 1:1000 dilution, Rockland Immunochemicals) and Alexa Fluor 488-conjugated Goat anti-rabbit antibody (1:1000 dilution, Life Technologies), and Anti-mCherry (mouse monoclonal, 1:1000 dilution Invitrogen) and HRP-conjugated Goat anti-mouse antibody (1:1000 dilution, Invitrogen). WesternBright ECL HRP substrate (Advansta) was used to detect the protein. Images were acquired using Image Lab software from Bio-Rad.

### Mass Spectrometry

For each sample, an equal volume of peptide was loaded onto a disposable Evotip C18 trap column (Evosep Biosystems, Denmark) as per the manufacturer’s instructions. Briefly, Evotips were wetted with 2-propanol, equilibrated with 0.1% formic acid, and then loaded using centrifugal force at 1200 g. Evotips were subsequently washed with 0.1% formic acid, and then 200 μL of 0.1% formic acid was added to each tip to prevent drying. The tipped samples were subjected to nanoLC on an Evosep One instrument (Evosep Biosystems). Tips were eluted directly onto a PepSep analytical column, dimensions: 8cm x 150µm, C18 column with 1.5 μm particle size (PepSep, Denmark), and a ZDV spray emitter (Bruker Daltronics). Mobile phases A and B were water with 0.1% formic acid (v/v) and 80/20/0.1% ACN/water/formic acid (v/v/v), respectively. The standard pre-set method of 60 samples-per-day was used, which is a 26 min gradient.

Mass spectrometry was performed on a hybrid trapped ion mobility spectrometry-quadrupole time of flight mass spectrometer (timsTOF Pro, Bruker Daltonics) with a modified nano-electrospray ion source (CaptiveSpray, Bruker Daltonics), as described previously (Shafraz et al., 2020). Mass spectrometry raw files were processed with MsFragger (Yu et al., 2020) as previously described (Shafraz et al., 2020). Volcano and PCA plots were generated using Spectronaut viewer. Known contamination proteins (Hodge et al., 2013) were removed, with further contamination filtering done by removing all non-canine proteins, with the exception of Cry/CIB from Arabidopsis (Ecad-LAB samples) and GFP from Aequorea (both), and filtering out all proteins categorized as “universal contaminants” by the software. A global background signal was added using the background signal imputation method, where the software chooses the best background signal to report as the precursor quantity. This allowed protein concentrations to be compared between the light positive and light negative conditions and ameliorated issues where proteins at too low concentrations in the negative condition could not be compared to the light positive condition. Only proteins that had two or more unique peptides detected were included in analysis. A Q-value of 0.05 was used to determine candidate threshold. Ecad-LAB runs and Ecad-Turbo runs were analyzed in separate files.

### Data Availability

MS raw spectra data can be found in the MassIVE database at: http://massive.ucsd.edu/ProteoSAFe/status.jsp?task=7dbf769e09fe410ba03980935ab8f048

## Acknowledgements

This research was supported in part by the National Institute of General Medical Sciences of the National Institutes of Health (R01GM121885) and the National Science Foundation (MCB-2022385). We thank Dr. Gabriela Grigorean for performing LC-MS/MS and data analysis in Proteomics Core Facility of the Genome Center at University of California, Davis.

## Author contributions

O.S., C.M.O.D., and S.S. designed research; O.S. and C.M.O.D. collected and analyzed data; O.S., C.M.O.D., and S.S. wrote the paper.

## Notes

### Competing Interest Statement

The authors have declared no competing interest.

### Summary of Updates

Results for additional imunofluorescence and mass spectrometry experiments described in the manuscript and SI. Figures 2 and 3 have been revised.

