## Supplementary Materials for "Light Activated BioID (LAB): an optically activated proximity labeling system to study protein-protein interactions"

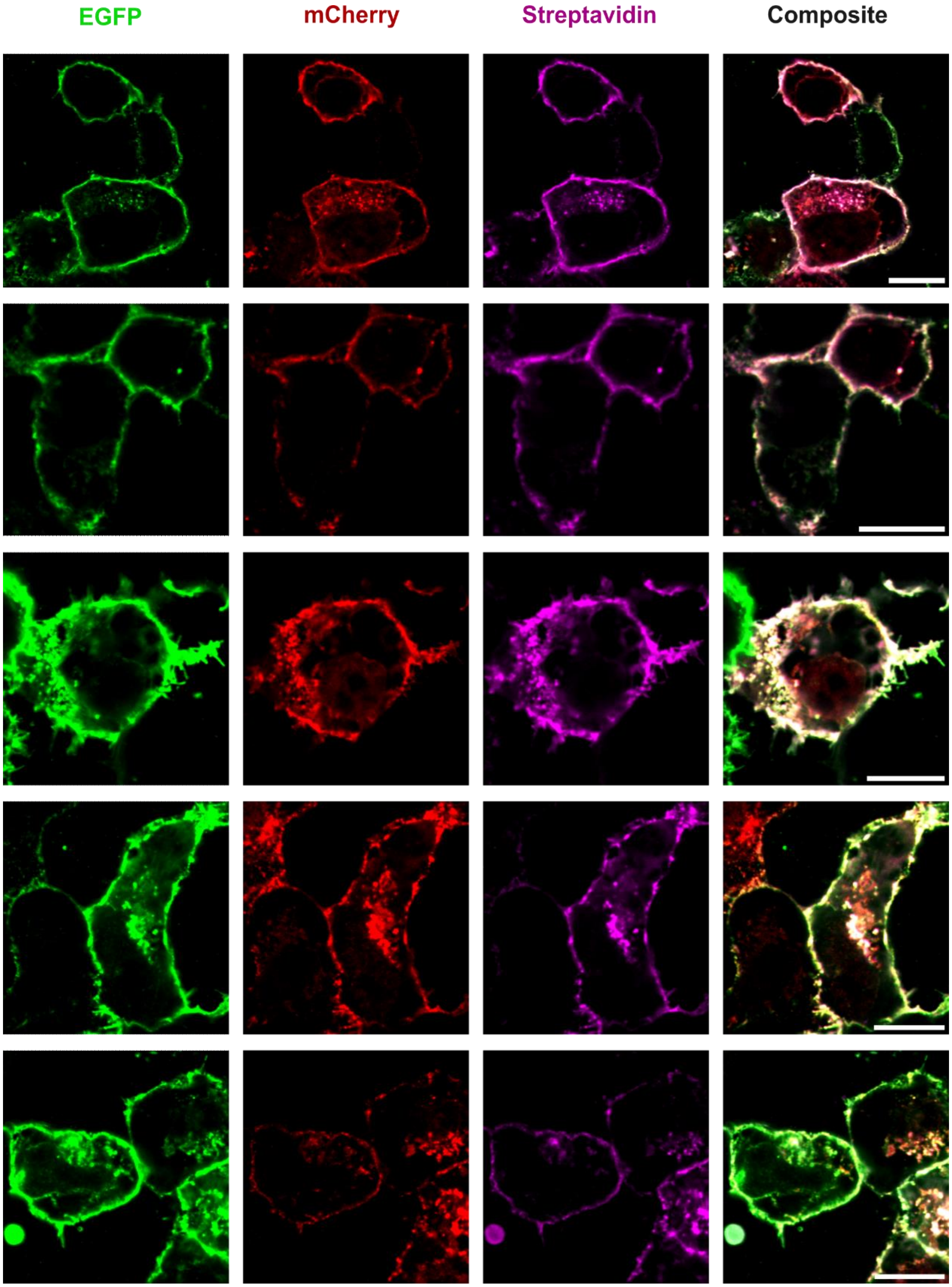


**Figure S1. LAB in HEK293T: Light [+] Biotin [+] condition**
Additional examples of LAB expressed in HEK293T cells exposed to light with added biotin as in Fig. 2 b. The composite images demonstrate robust CibN / CryC colocalization as well as strong biotinylation. All scale bars 10μm.

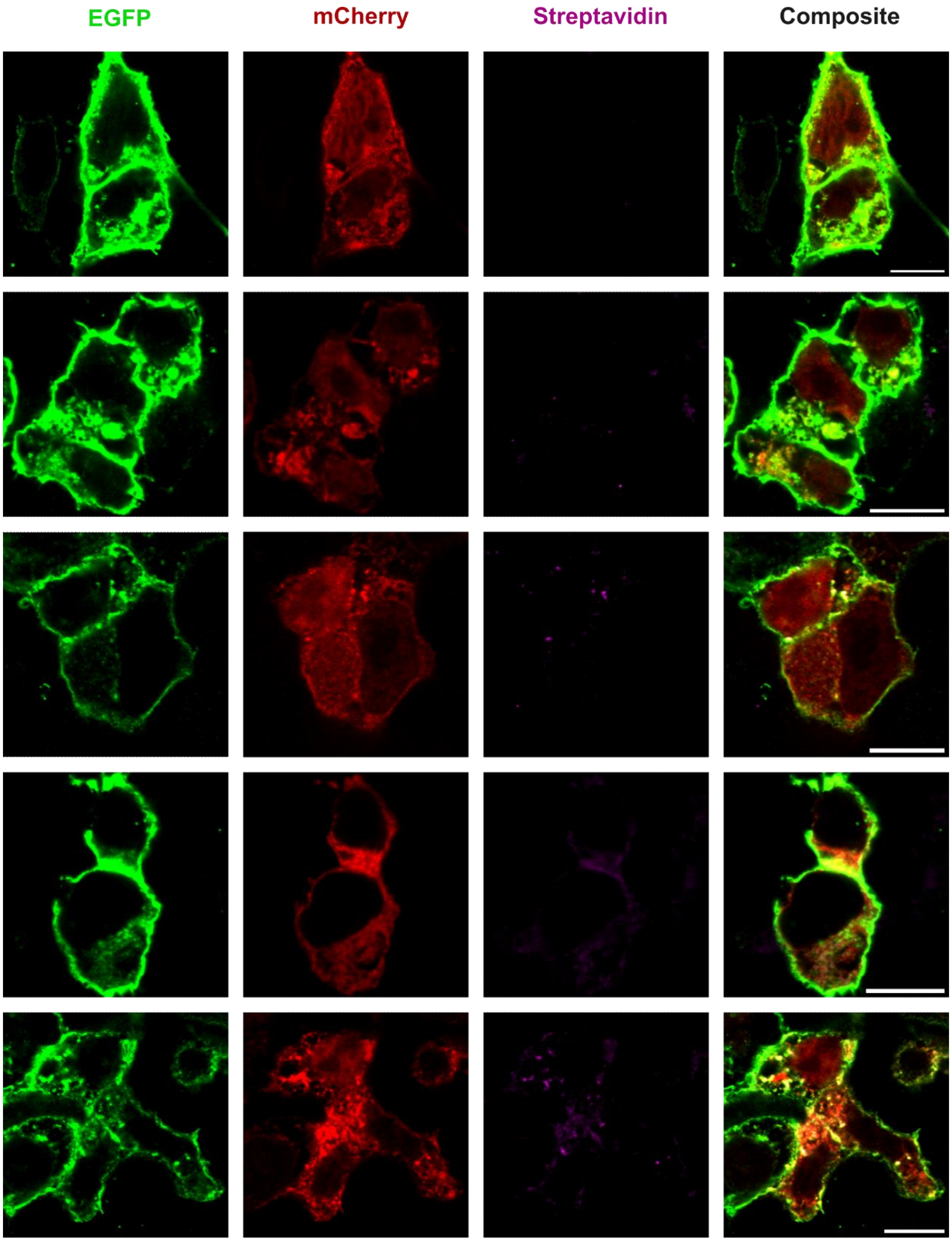


**Figure S2. LAB in HEK293T: Light [-] Biotin [+] condition**
Additional examples of LAB expressed in HEK293T cells incubated with added biotin in darkness as in Fig. 2 c. The composite images demonstrate insignificant CibN / CryC colocalization as well as a lack of biotinylation. All scale bars 10μm.

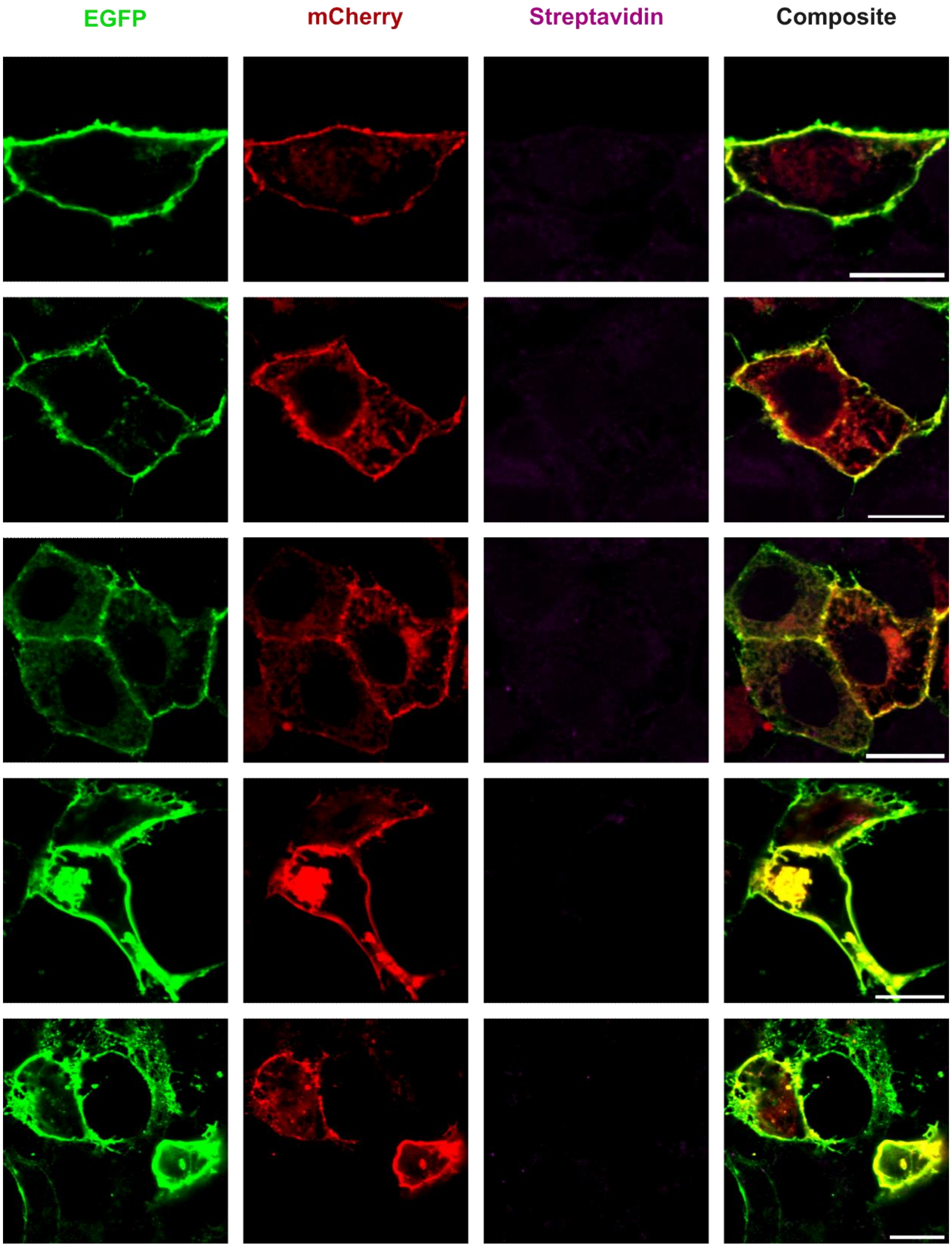


**Figure S3. LAB in HEK293T: Light [+] Biotin [-] condition**
Additional examples of LAB expressed in HEK293T cells exposed to light without added biotin as in Fig. 2 d. The composite images demonstrate the strong CibN / CryC colocalization but an absence of biotinylation. All scale bars 10μm.

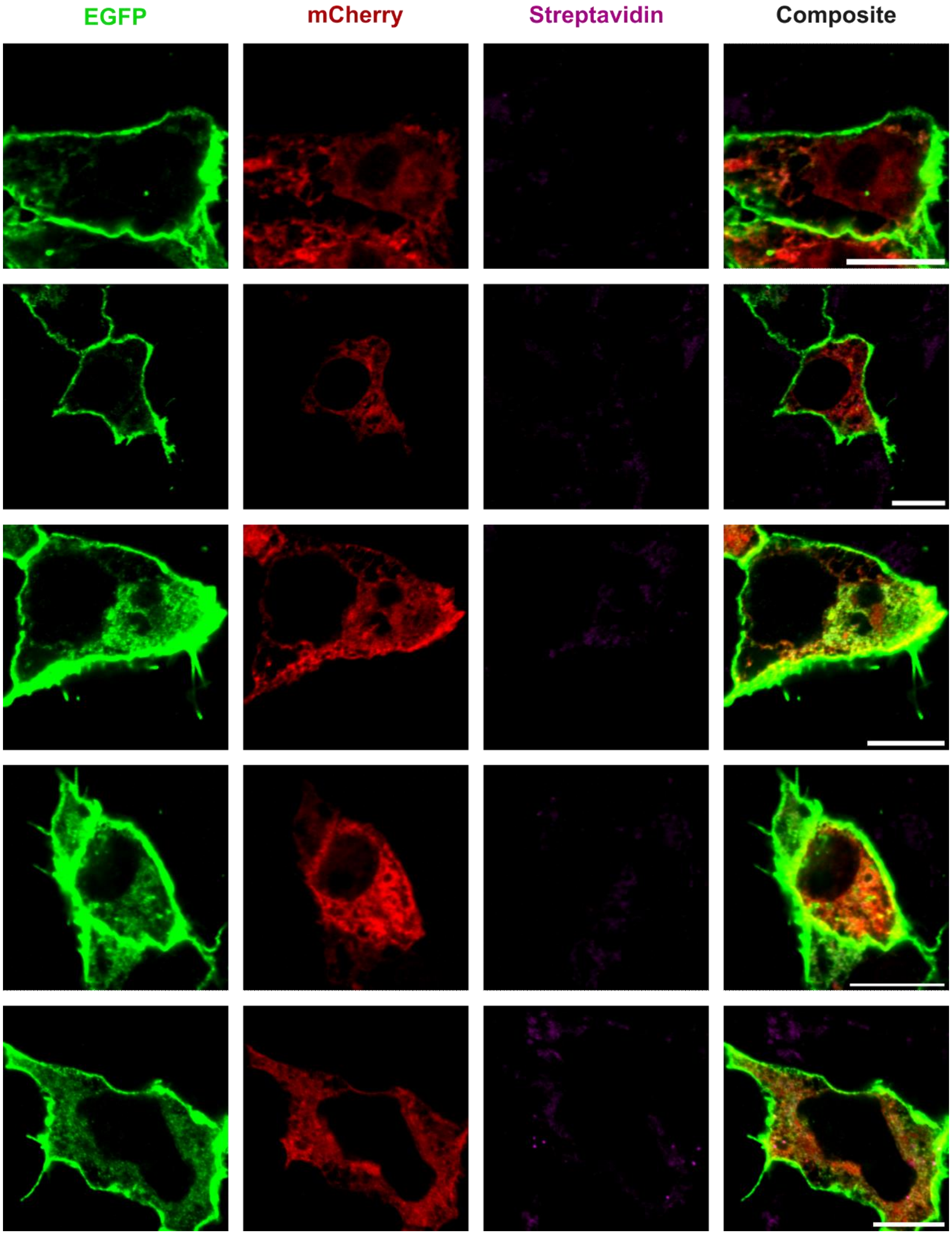


**Figure S4. LAB in HEK293T: Light [-] Biotin [-] condition**
Additional examples of LAB expressed in HEK293T cells in the absence of light and biotin as in Fig. 2 e. The composite images demonstrate insignificant CibN / CryC colocalization as well as a lack of biotinylation. All scale bars 10μm.

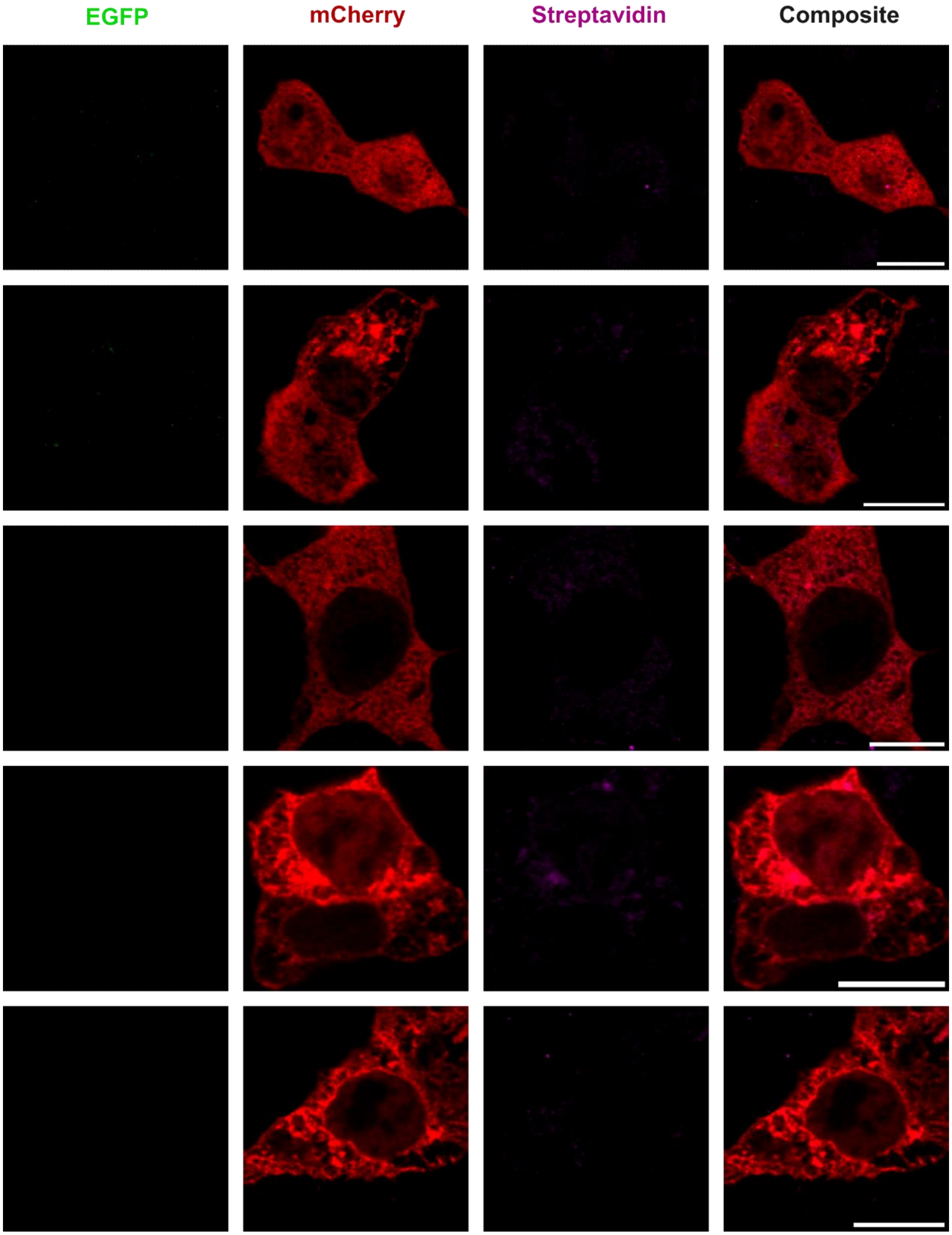


**Figure S5. HEK293T: CIB [-] Cry [+] Light [+] Biotin [+] condition**
Additional examples of HEK293T cells transfected with solely CryC in the presence of light and biotin as in Fig. 2 f. The composite images demonstrate no biotinylation. All scale bars 10μm.

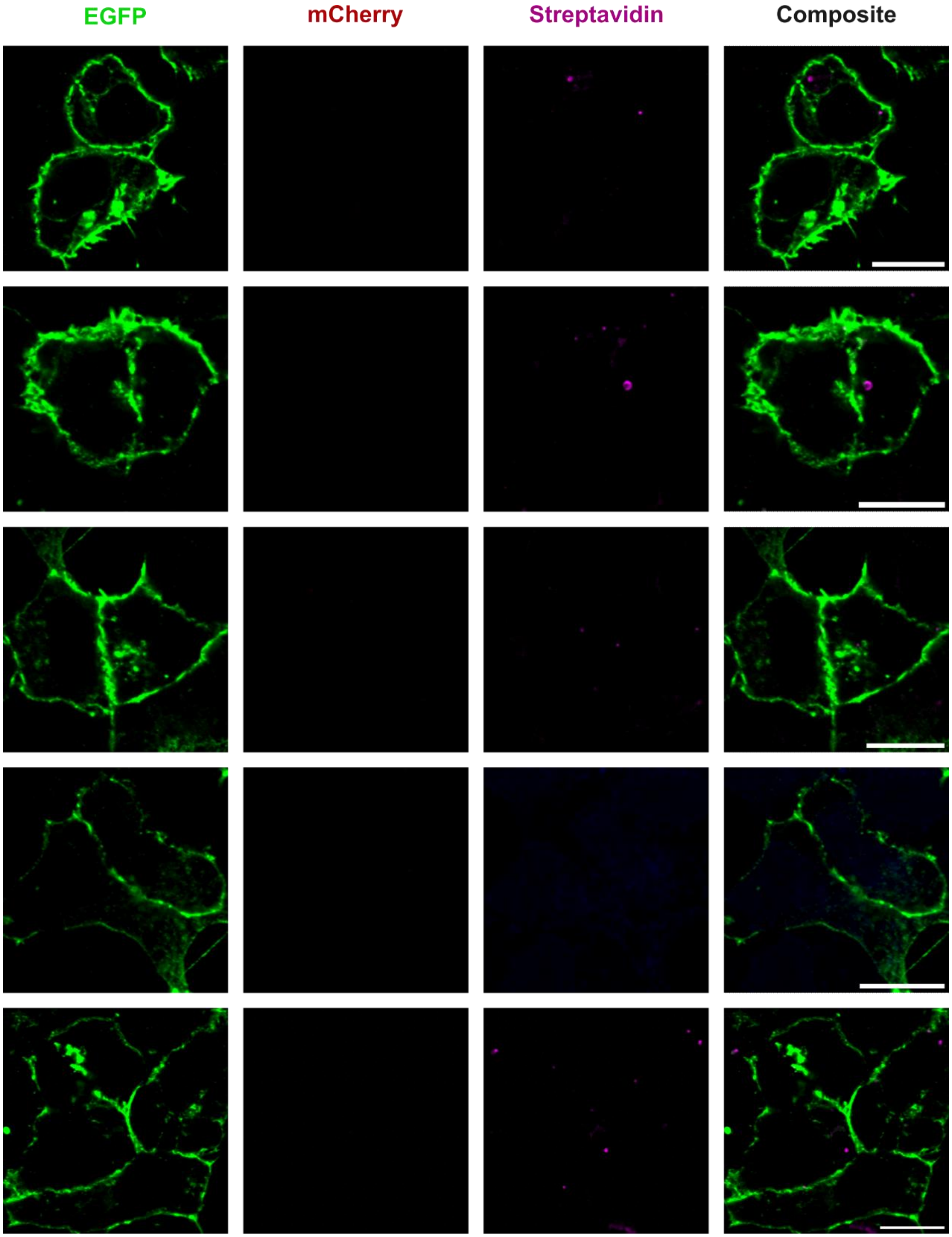


**Figure S6. HEK293T: CIB [+] Cry [-] Light [+] Biotin [+] condition**
Additional examples of HEK293T cells transfected only with CibN in the presence of light and biotin as in Fig. 2 g. The composite images demonstrate a lack of biotinylation. All scale bars 10μm.

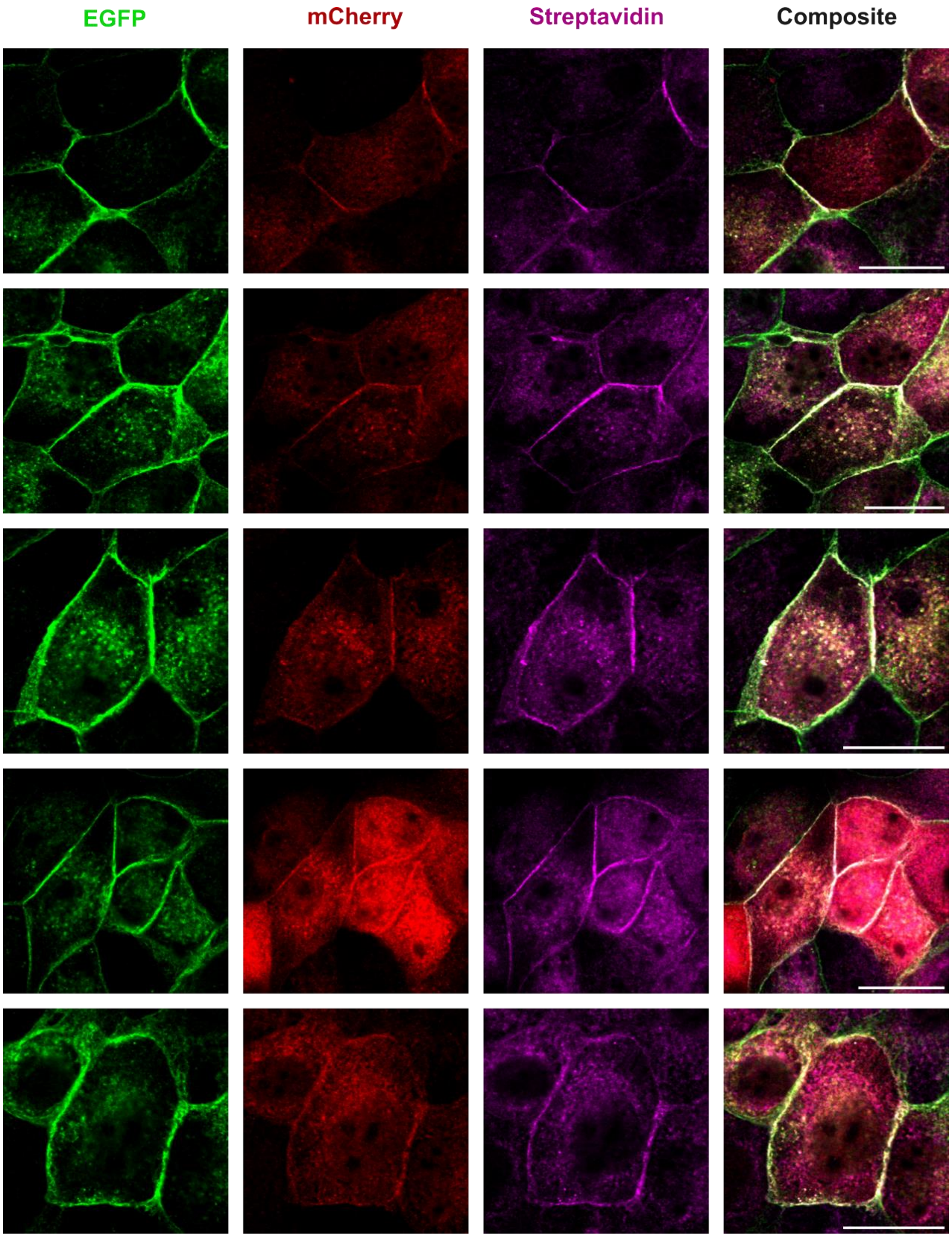


**Figure S7. Ecad-LAB in MDCK: Light [+] Biotin [+] condition**
Additional examples of Ecad-LAB stably expressed in MDCK cells exposed to light with added biotin as in Fig. 3 b. The composite images demonstrate robust ECibN / CryC colocalization as well as strong biotinylation. All scale bars 20μm.

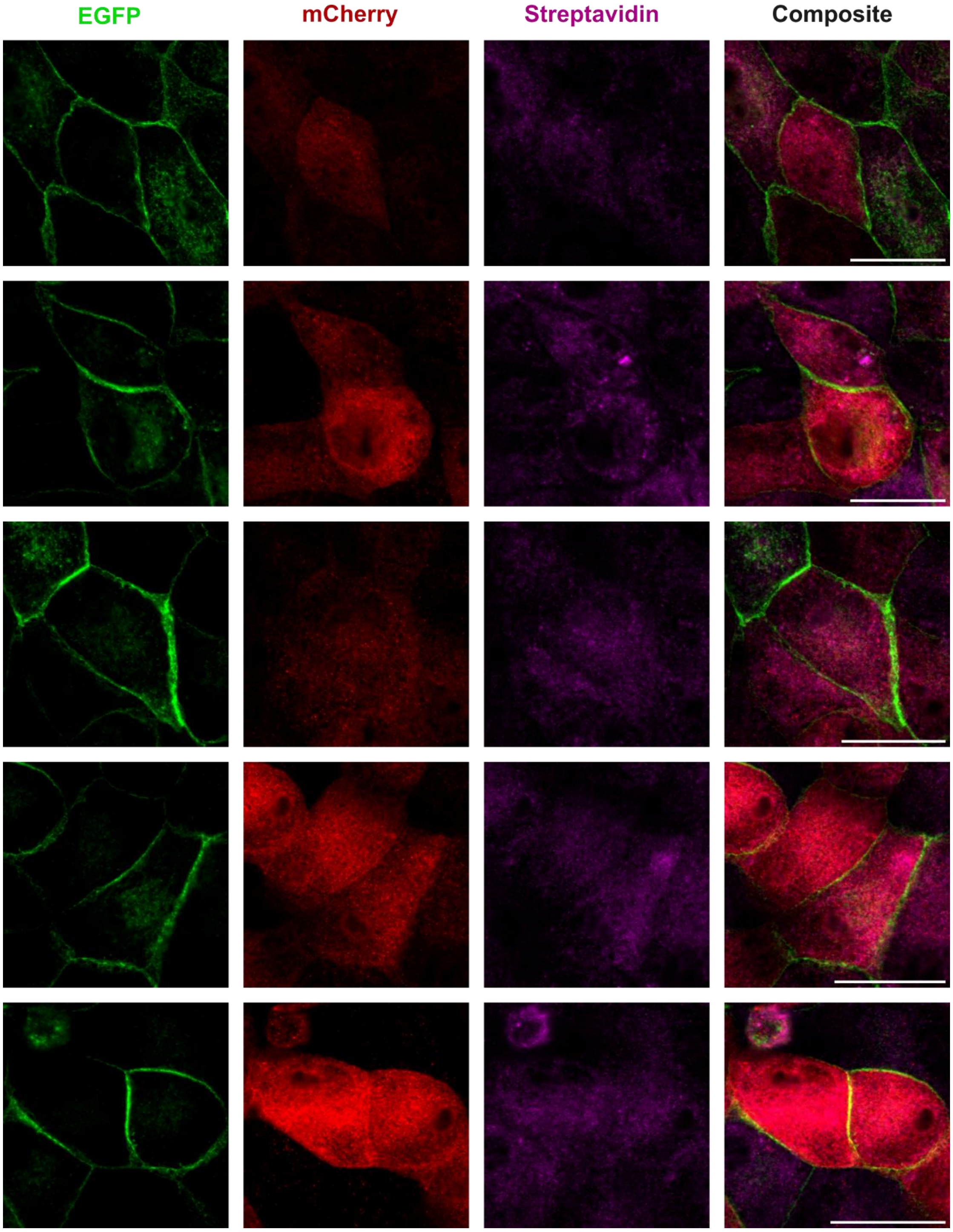


**Figure S8. Ecad-LAB in MDCK: Light [-] Biotin [+] condition**
Additional examples of Ecad-LAB stably expressed in MDCK cells incubated with added biotin in darkness as in Fig. 3 c. The composite images demonstrate insignificant ECibN / CryC colocalization as well as a lack of biotinylation. All scale bars 20μm.

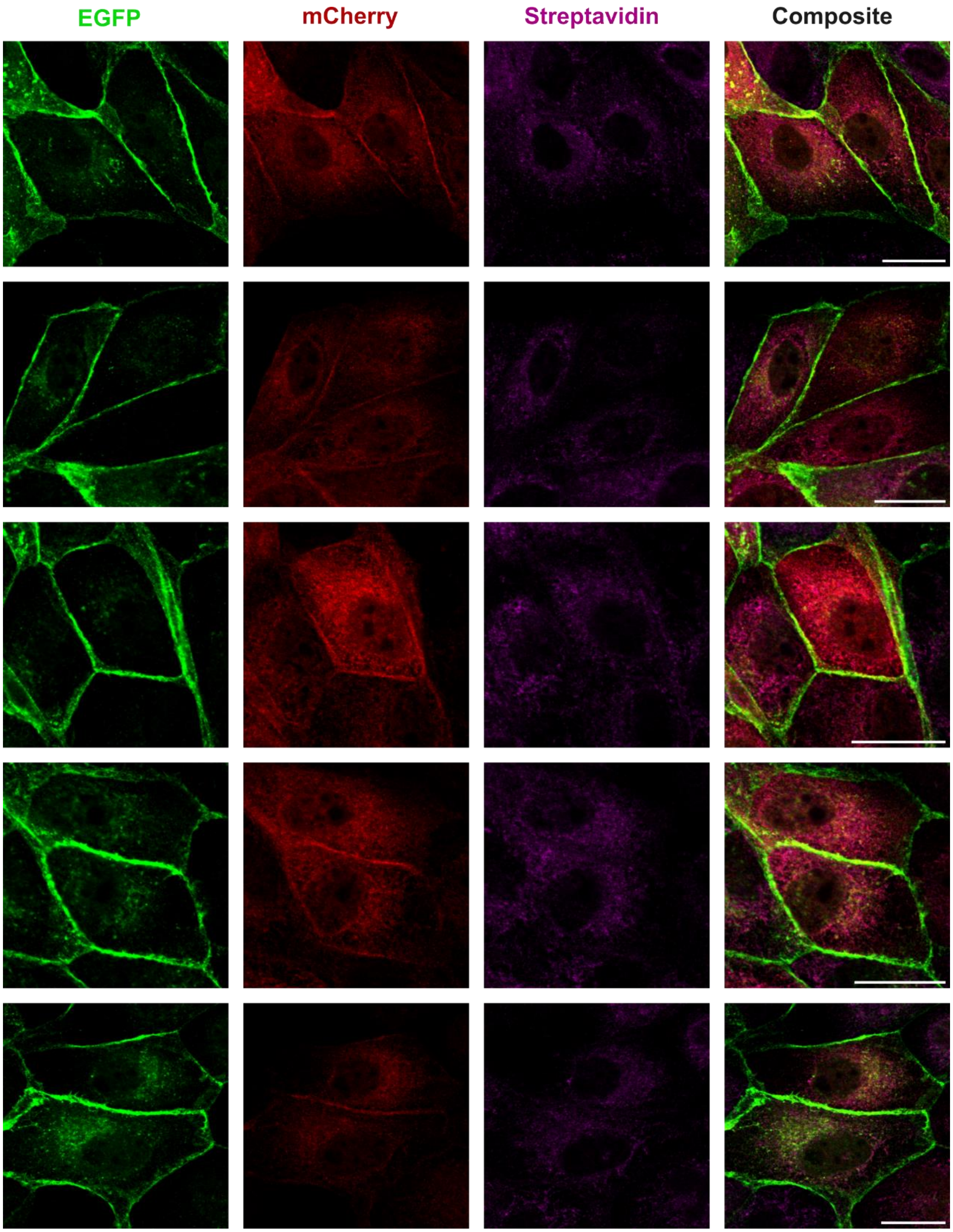


**Figue S9. Ecad-LAB in MDCK: Light [+] Biotin [-] condition**
Additional examples of Ecad-LAB stably expressed in MDCK cells exposed to light without added biotin as in Fig. 3 d. The composite images demonstrate strong ECibN / CryC colocalization but an absence of biotinylation. All scale bars 20μm.

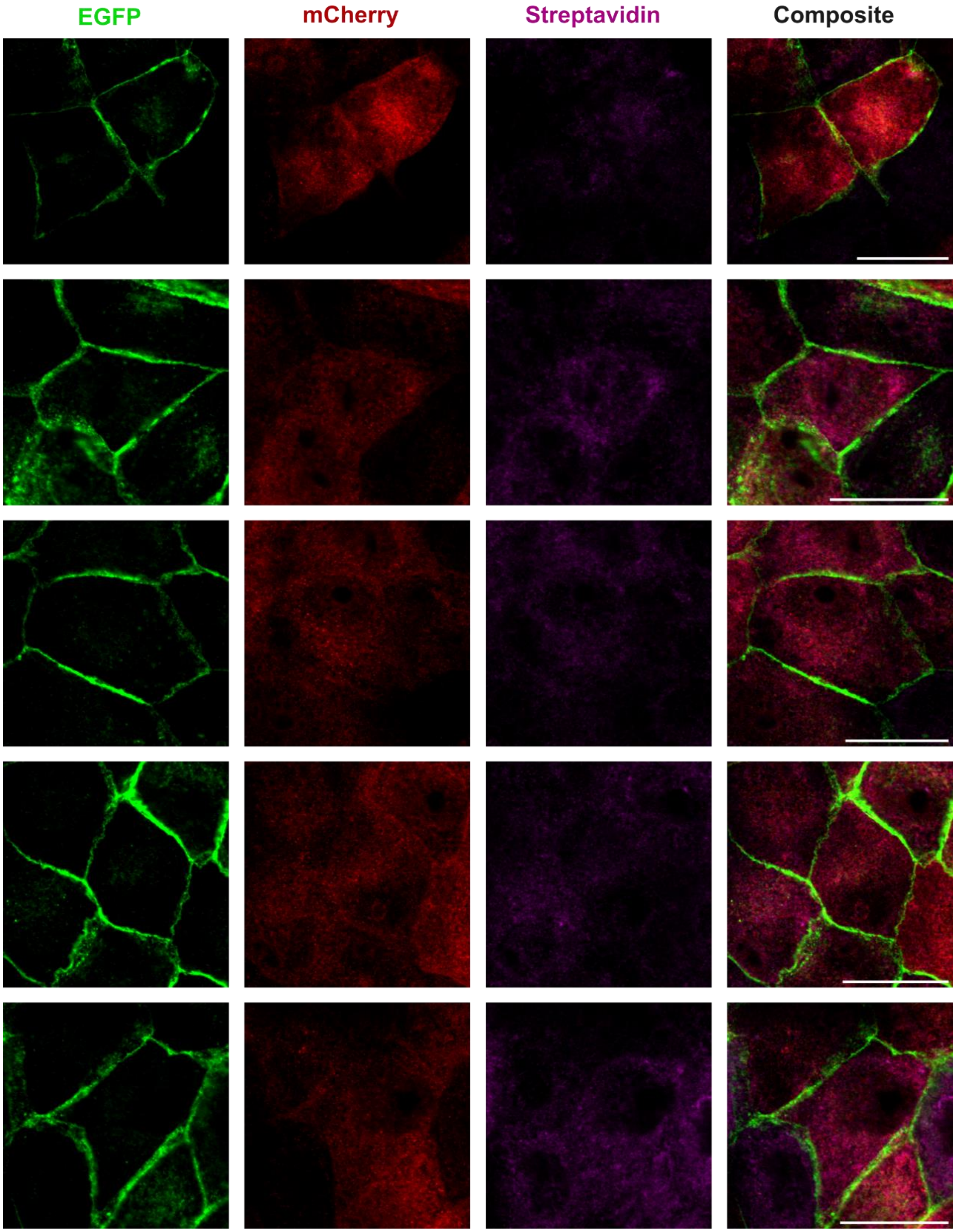


**Figure S10. Ecad-LAB in MDCK: Light [-] Biotin [-] condition**
Additional examples of Ecad-LAB stably expressed in MDCK cells in the light and biotin negative condition as in Fig. 3 e. The composite images demonstrate insignificant ECibN / CryC colocalization as well as a lack of biotinylation. All scale bars 20μm.

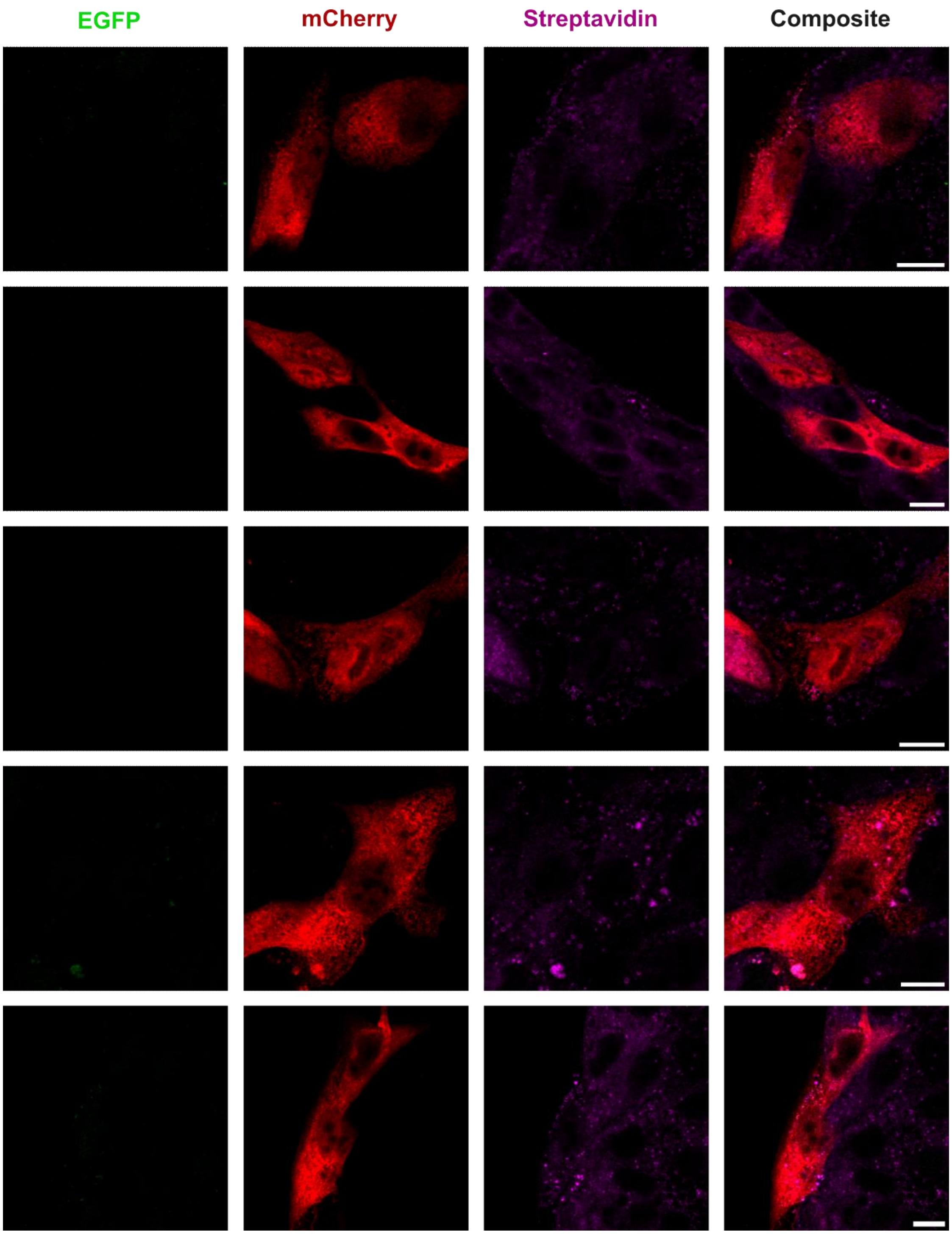


**Figur S11. MDCK: CIB [-] Cry [+] Light [+] Biotin [+] condition**
Additional examples of MDCK cells transiently transfected with solely CryC in the presence of light and biotin as in Fig. 3 f. The composite images demonstrate no ECibN / CryC colocalization as well as a lack of biotinylation. All scale bars 10µm.

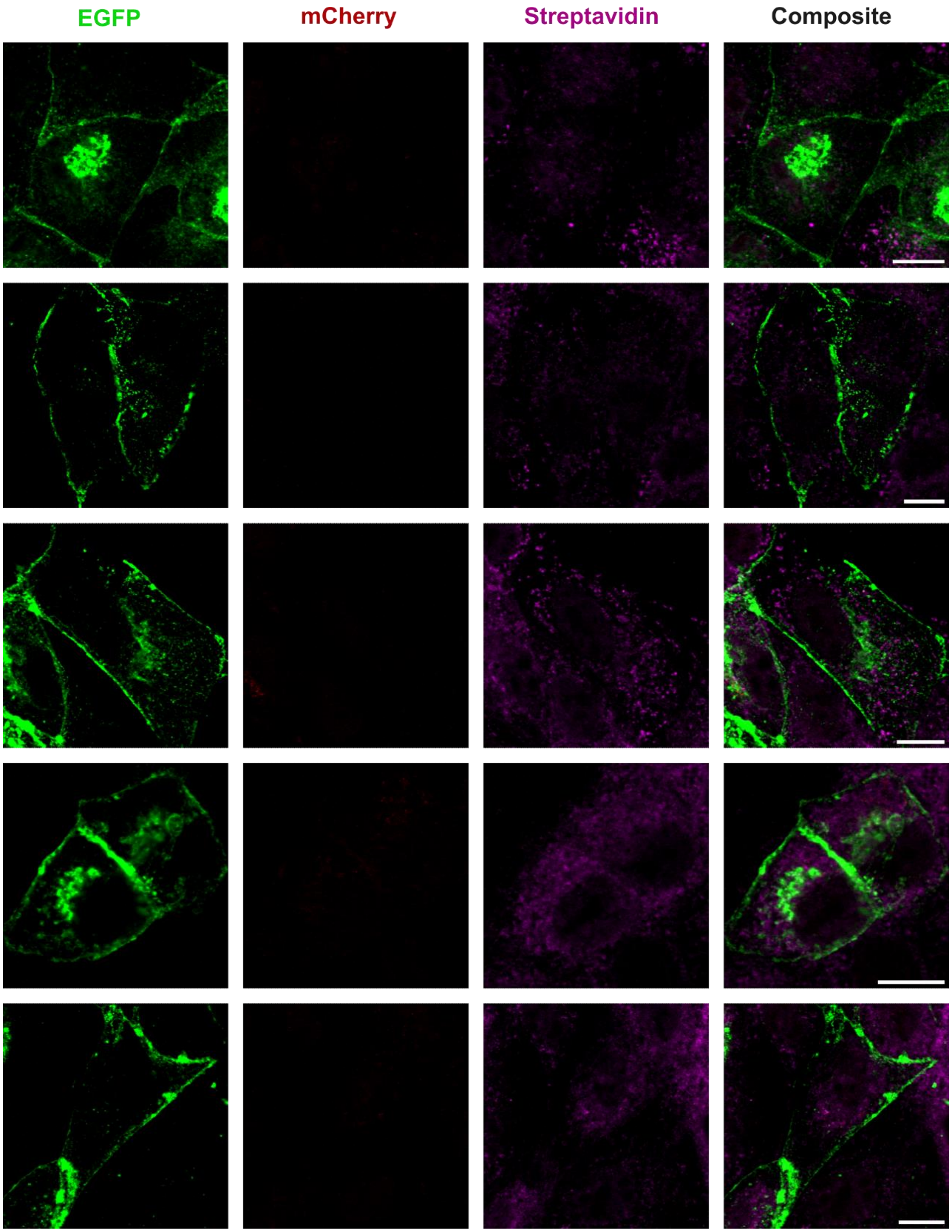


**Figure S12. MDCK: CIB [+] Cry [-] Light [+] Biotin [+] condition**
Additional examples of MDCK cells transiently transfected with solely ECibN in the presence of light and biotin as in Fig. 3 g. The composite images demonstrate no ECibN / CryC colocalization as well as a lack of biotinylation. All scale bars 10μm.

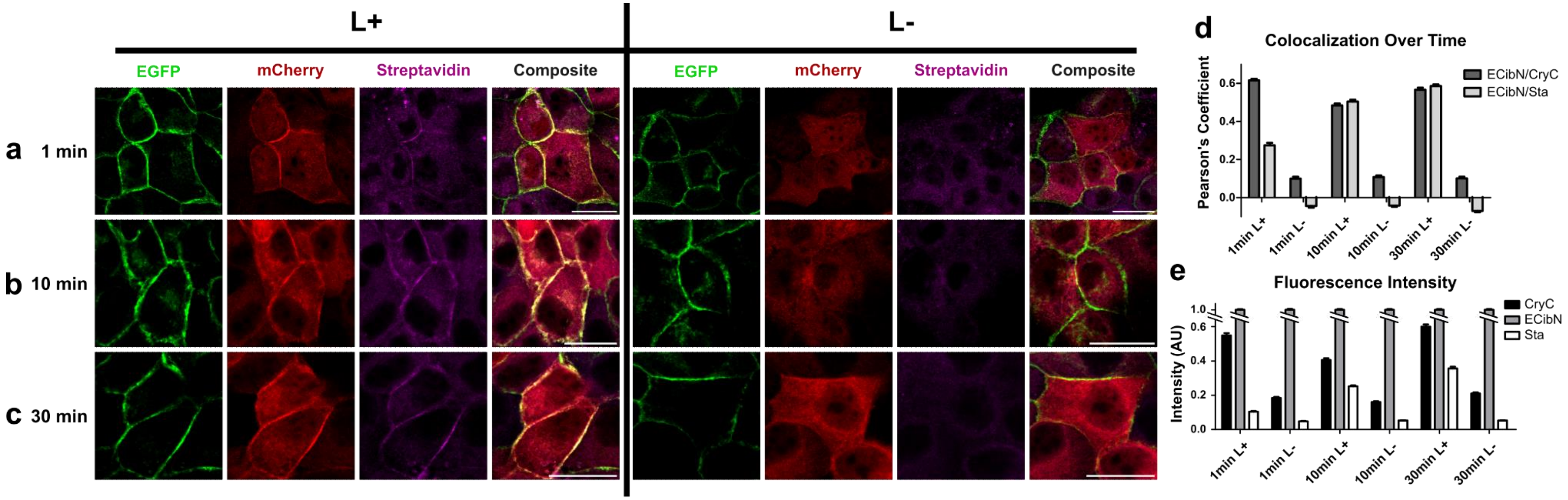


**Figure S13. Biotinylation time dependence.**

Immunofluorescence images of Ecad-LAB cells incubated with 100µM biotin, after exposure to light for (a) 1 min, (b) 10 min, and (c) 30 min. Cells were stained for GFP (green), mCherry (red), and streptavidin (magenta). Light positive (L+) images are on the left, while light negative (L-) images are on the right. The panels in each column display identical minimum and maximum intensities. Scale bars are 20µm. For all measured light exposures, membrane localization of Cry and biotinylation was detected compared to the L- condition. d) Average Pearson's coefficient shows that ECibN and CryC colocalize in the presence of light at all time points, while Sta colocalizes to ECibN with increasing efficiency over time. e) Fluorescence intensity of CryC, ECibN, and Sta, normalized to ECibN intensity to account for variation in expression levels in different cells, shows that CryC only associates to the membrane in light. Biotinylation is detected above background after only one minute and continues to increase in intensity over time. P-values for Sta signal between L+ and L- conditions are as follows: 1min: 4.07E-29; 10min: 1.69E-91; 30min: 6.13E-78. Errors: s.e., n's are as follows: 1min L+: 169; 1min L-: 137; 10min L+: 191; 10min L-: 222; 30min L+: 204; 30min L-: 192, split between three biological replicates for all conditions.

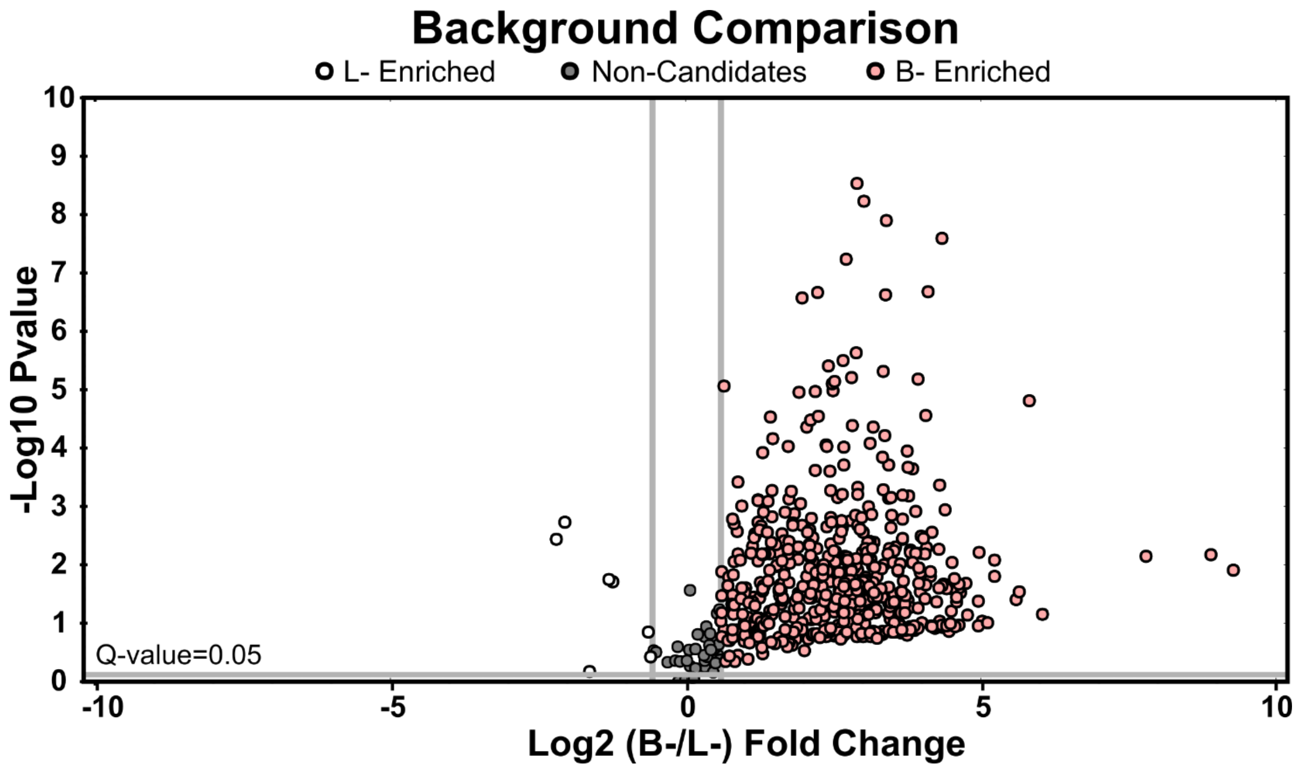


**Figure S14. Ecad-Turbo and Ecad-LAB Background.** Volcano plot showing the relationship between the detected levels of protein in the Ecad-Turbo biotin negative condition (B-; pink) and the Ecad-LAB light negative condition (L-; white).

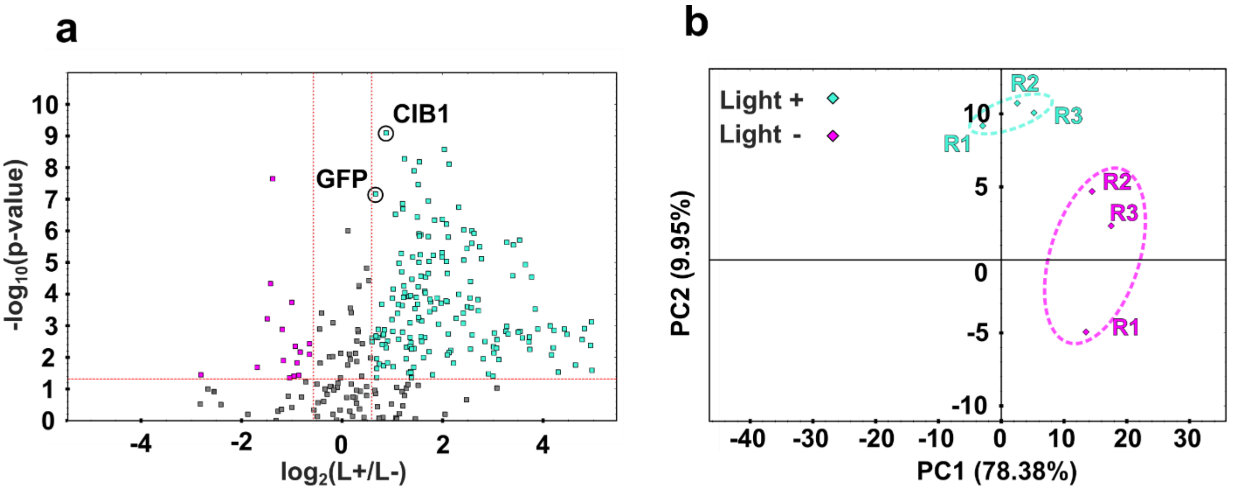


**Figure S15: Mass spectrometry (MS) analysis for LAB in HEK293T cells.** (a) MS data for HEK293T cells show more biotinylated proteins for light-exposed cells (cyan) compared to cells kept in dark (magenta), with CIB1 and GFP (circled) being among the top-ranked in the data. (b) Principal Component Analysis (PCA) shows similar variation for all replicates of the same condition (R1, R2, R3), with light-exposed cells (cyan) calculated to be significantly different from the control cells kept in the dark (magenta) with 100 µM added biotin.

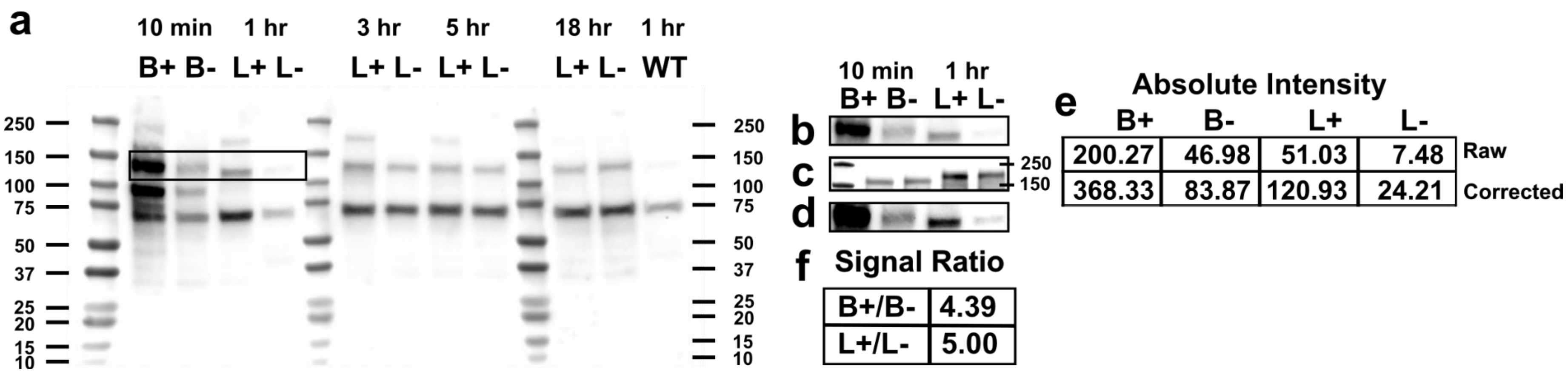


**Figure S16. Western Blots comparing Ecad-LAB and Ecad-Turbo biotinylation efficiency.** (a) Western Blots comparing biotinylation efficiencies of Ecad-Turbo and Ecad-LAB expressed in stabilized WT MDCK cells for different time points. Ecad-Turbo was exposed to either the presence (B+) or absence (B-) of 100µM biotin for 10 minutes. Ecad-LAB was incubated in 100µM biotin, in the presence (L+) or absence (L-) of light for 1 hr, 3 hrs, 5 hrs, and 18 hrs, respectively. Whole lysate was stained with anti-biotin antibody. Final lane (WT) is lysate from un-transfected cells. The same amount of total protein was loaded into each lane. While Ecad-Turbo has a markedly higher biotinylation efficiency than Ecad-LAB, the larger number of false positives in Ecad-Turbo (see Fig. 4 and S14) coupled with the ~44% localization of Cry and CIB (see Fig. 3) makes a direct comparison of biotinylation efficiencies difficult. We therefore compared the biotinylation efficiency of a representative band from the first four lanes (marked by the square box). (b) Blown up view of highlighted bands used in subsequent analysis comparing Ecad-Turbo after 10min of B(+) or B(-) with Ecad-LAB after 1hr of L(+) or L(-). (c) Western Blot stained with Anti-Ecad showing Ecad-Turbo (Lanes 2- 3) or ECibN (Lanes 4-5) in order to compare construct expression levels in stabilized WT MDCK cells. (d) Band intensity (in *b*) corrected for construct expression using Ecad expression levels (panel *c*) and Pearson's coefficient. (e) Raw and corrected intensities of the bands, measured using imageJ. (f) The ratio between the positive and negative condition bands for each construct shows that Ecad-LAB exposed to light for 1 hour has a higher biotinylation efficiency than Ecad-Turbo incubated with biotin for 10 minutes. L+ cells were exposed to an altered light cycle of 1min on, 5min off to reduce phototoxicity in longer time conditions.

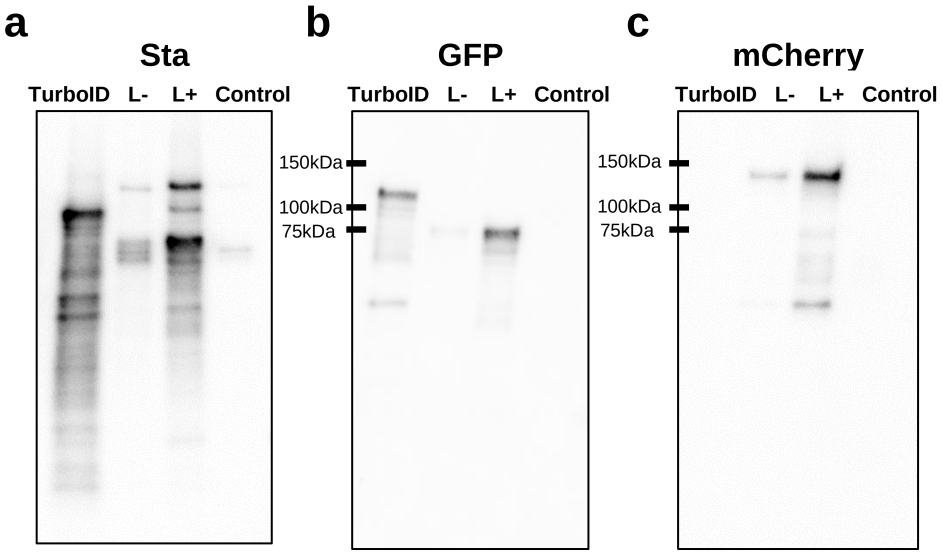


**Figure S17. Western blots detect biotinylation, CibN and CryC in HEK cells.** Western blots run using protein harvested from cells transiently transfected with CIB1-TurboID-pmEGFP (TurboID), LAB, and blank HEK293T as a control. Biotinylated proteins were pulled down on Sta conjugated beads, eluted, and then stained for (a) Sta, (b) GFP, or (c) mCherry. All samples were incubated with 100μM biotin for one hour, and the L+ condition was exposed to blue light on alternating 10 min intervals for 1hr. L+ lanes in all gels show a significant increase in biotinylated protein compared to the L- lanes. The L+ lane in (a) also shows significantly less biotinylation than the full-length TurboID lane. The Control lane in (a) shows a small band of endogenously biotinylated proteins. (b) As GFP is conjugated to both TurboID and CibN, the TurboID and LAB lanes correctly show GFP presence while the Control lane lacks it. (c) As mCherry is conjugated to CryC and is not present on the full-length TurboID, mCherry correctly shows up in only the LAB lanes and neither the TurboID or Control lanes.

| Normalized Fluorescent Intensity ± s.e. (AU) | | |
|---|---|---|
| L+ B+ | CibN | 1.000 ± 0.037 |
| | CryC | 0.556 ± 0.044 |
| | Sta | 0.69 ± 0.047 |
| L- B+ | CibN | 1.000 ± 0.043 |
| | CryC | 0.114 ± 0.012 |
| | Sta | 0.048 ± 0.005 |
| L+ B- | CibN | 1.000 ± 0.048 |
| | CryC | 0.402 ± 0.047 |
| | Sta | 0.016 ± 0.001 |
| L- B- | CibN | 1.000 ± 0.047 |
| | CryC | 0.201 ± 0.025 |
| | Sta | 0.017 ± 0.002 |

**Table S1. Normalized fluorescent intensity for HEK cells**

| Pearson's Coefficient ± s.e. | | |
|---|---|---|
| L+ B+ | CibN / CryC | 0.59 ± 0.02 |
| | CibN / Sta | 0.57 ± 0.02 |
| L- B+ | CibN / CryC | 0.15 ± 0.03 |
| | CibN / Sta | 0.17 ± 0.03 |
| L+ B- | CibN / CryC | 0.59 ± 0.02 |
| | CibN / Sta | 0.09 ± 0.02 |
| L- B- | CibN / CryC | 0.17 ± 0.03 |
| | CibN / CryC | -0.00 ± 0.01 |

**Table S2. Pearson's Coefficient for HEK cells**

| Fluorescence P Values (Two-tailed T Test) | | |
|---|---|---|
| L+B+/ L-B+ | CryC / CryC | 9.08E-13 |
| | Sta / Sta | 7.78E-17 |
| L+B+/ L+B- | CryC / CryC | 0.004 |
| | Sta / Sta | 3.66E-13 |
| L+B+/ L-B- | CryC / CryC | 1.12E-08 |
| | Sta / Sta | 5.70E-17 |

**Table S3. P Values for HEK cell Fluorescence**

| Pearson's Coefficient P Values (Two-tailed T Test) | | |
|---|---|---|
| L+B+/ L-B+ | CibN / CryC | 2.62E-15 |
| | CibN / Sta | 4.55E-13 |
| L+B+/ L+B- | CibN / CryC | 0.77 |
| | CibN / Sta | 1.35E-13 |
| L+B+/ L-B- | CibN / CryC | 3.35E-11 |
| | CibN / Sta | 2.83E-20 |

**Table S4. P Values for HEK cell Pearson's Coefficients**

| Normalized Fluorescent Intensity ± s.e. (AU) | | |
|---|---|---|
| L+B+ | ECibN | 1.000 ± 0.011 |
| | CryC | 0.174 ± 0.005 |
| | Sta | 0.410 ± 0.012 |
| L-B+ | ECibN | 1.000 ± 0.015 |
| | CryC | 0.092 ± 0.003 |
| | Sta | 0.054 ± 0.002 |
| L+B- | ECibN | 1.000 ± 0.012 |
| | CryC | 0.159 ± 0.005 |
| | Sta | 0.050 ± 0.002 |
| L-B- | ECibN | 1.000 ± 0.014 |
| | CryC | 0.074 ± 0.002 |
| | Sta | 0.052 ± 0.002 |

**Table S5. Normalized fluorescent intensity for MDCK cells**

| Pearson's Coefficient | | |
|---|---|---|
| L+B+ | CibN / CryC | 0.44 ± 0.03 |
| | CibN / Sta | 0.61 ± 0.02 |
| L-B+ | CibN / CryC | 0.08 ± 0.02 |
| | CibN / Sta | 0.00 ± 0.01 |
| L+B- | CibN / CryC | 0.41 ± 0.01 |
| | CibN / Sta | 0.03 ± 0.02 |
| L-B- | CibN / CryC | 0.12 ± 0.01 |
| | CibN / Sta | -0.01 ± 0.01 |

**Table S6. Pearson's Coefficient for MDCK cells**

| Fluorescence P Values (Two-tailed T Test) | | |
|---|---|---|
| L+B+/ L-B+ | CryC/CryC | 7.86E-35 |
| | Sta / Sta | 3.41E-67 |
| L+B+/ L+B- | CryC/CryC | 0.03 |
| | Sta / Sta | 2.83E-67 |
| L+B+/ L-B- | CryC/CryC | 8.33E-43 |
| | Sta / Sta | 3.60E-69 |

**Table S7. P Values for MDCK cell Fluorescence**

| Pearson's Coefficient P Values (Two-tailed T Test) | | |
|---|---|---|
| L+B+/ L-B+ | ECibN / CryC | 1.22E-64 |
| | ECibN / Sta | 2.40E-112 |
| L+B+/ L+B- | ECibN / CryC | 0.01 |
| | ECibN / Sta | 3.21E-107 |
| L+B+/ L-B- | ECibN / CryC | 3.23E-56 |
| | ECibN / Sta | 4.26E-111 |

**Table S8. P Values for MDCK cell Pearson's Coefficients**

| | | |
|---|---|---|
| CIB1 | Forward (F) | 5’-ttagtgaaccgtcagatccgctagcccATGAATGGAGCTATAGGAG |
| | Reverse (R) | 5’-tgccgatatcTACTCCTAAATTGCCATAGAG |
| spTurboIDN | F | 5’-tttaggagtagatatcGGCAAGCCCATCCCCAAC |
| | R | 5’-cttgctcaccatggtggcgaccggtccactcccCAGAATCTGTTTAGCGTTCAGCAG |
| spTurboIDC | F | 5’-ggaaaaaatggttgcaaagcccggggtAAGGGCTCGGGCTCGACC |
| | R | 5’-atggtggcgaccggtggatcgcccgagcccttCTTTTCGGCAGACCGCAGAC |
| EGFP | F | 5’-tggaggtggcgaggacgacctcgagATGGTGAGCAAGGGCGAG |
| | R | 5’-tgattatgatctagagtcgcggccgcttagaagcttgaCTTGTACAGCTCGTCCATGC |
| V5_spTN | F | 5’-ttagtgaaccgtcagatccgctagcatgGGCAAGCCCATCCCCAAC |
| | R | 5’-tagctccattcatggtggcgcccgagcccttCAGAATCTGTTTAGCGTTCAGCAG |
| spTNCIB | F | 5’-tggacgagctgtacaagtcaagcttcGGCAAGCCCATCCCCAAC |
| | R | 5’-tgattatgatctagagtcgcggccgcttaTACTCCTAAATTGCCATAGAGATTCTGC |

**Table S9. List of primers used for PCR amplification**
